# Convergent evolution of the Hedgehog/Intein fold in protein splicing

**DOI:** 10.1101/2020.03.19.998260

**Authors:** Hannes M. Beyer, Salla I. Virtanen, A. Sesilja Aranko, Kornelia M. Mikula, George T. Lountos, Alexander Wlodawer, O. H. Samuli Ollila, Hideo Iwaï

**Affiliations:** Institute of Biotechnology, University of Helsinki. P.O. Box 65, Helsinki, FIN-00014, Finland; Basic Science Program, Frederick National Laboratory for Cancer Research, Frederick, MD 21702, USA; Macromolecular Crystallography Laboratory, National Cancer Institute, Frederick, MD 21702, USA

**Keywords:** protein-splicing mechanism, intein, evolution, protease

## Abstract

The widely used molecular evolutionary clock assumes the divergent evolution of proteins. Convergent evolution has been proposed only for small protein elements but not for an entire protein fold. We investigated the structural basis of the protein splicing mechanism by class 3 inteins, which is distinct from class 1 and 2 inteins. We gathered structural and mechanistic evidence supporting the notion that the Hedgehog/INTein (HINT) superfamily fold, commonly found in protein splicing and related phenomena, could be an example of convergent evolution of an entire protein fold. We propose that the HINT fold is a structural and biochemical solution for *trans*-peptidyl and *trans*-esterification reactions.

## Introduction

Proteins fold into various defined three-dimensional structures to exert their unique biochemical functions. Proteins with similar structures and functions across different organisms share common ancestors and have evolved through divergent evolution^1^. However, protein structures could also converge into a similar structure to function analogously but having evolved from different ancestors. This convergent evolution is best exemplified by the catalytic Ser-His-Asp triad commonly found in hydrolases, suggesting the importance of structural and functional constraints required for catalysis^2,3,4^. Even though convergent evolution is a commonly observed phenomenon across the diversity of living organisms, the convergent evolution of protein structures has been documented for only small structural elements of proteins^5^. Structural convergence of an entire protein fold has not been reported^6^.

Protein splicing catalyzed by intervening protein sequences termed inteins was discovered in the 1990s. The splicing reaction involves the self-removal of the intein and concomitant joining of the two flanking sequences (exteins)^7,8^. Protein splicing is analogous to RNA splicing but occurs on the protein level. The biological function of protein splicing is still enigmatic despite several proposals for eventual regulatory functions^9^. Inteins are often considered merely as selfish gene elements because they can be removed without affecting the fitness of their host organisms. Inteins commonly insert in conserved sequences close to the active sites of essential proteins. Any mutations within inteins detrimental to the protein splicing could be lethal or strongly affect the fitness of their host, which is the mechanism ensuring intein persistence and protection from degeneration.

The most common protein splicing mechanism has been generally accepted and involves the four concerted steps: (1) N-S(O) acyl shift, (2) *trans*-(thio)esterification (esterification), (3) Asn cyclization, and (4) S(O)-N acyl shift (Fig. 1a)^10^. Inteins undergoing the canonical splicing mechanism are referred to as class 1 inteins (Fig. 1a)^11^. Not all of the four steps are exploited among all Hedgehog/INTein (HINT) superfamily members, which all share the same flat horseshoe-like HINT fold and catalyze protein splicing as well as related reactions (Fig. 1b)^8,12^. For example, the C-terminal domain of the Hedgehog protein (Hh-C or hog domain), a member of the HINT superfamily, uses the first step of the N-S acyl shift for cholesterol modification of the N-terminal signaling domain (Hh-N)^8,12^. Bacterial Intein-Like (BIL) domains lack the nucleophilic +1 residue of inteins essential for the *trans*-esterification step in the protein-splicing reaction and produce predominantly cleaved products^13^. Some inteins do not undergo the canonical splicing reaction of class 1 inteins. Inteins without the first nucleophilic residue required for the initial N-S(O) acyl shift step were originally termed class 2 (Fig. 1a)^14^. However, class 3 inteins lacking the N-terminal serine or cysteine, similar to class 2 inteins, have been identified. Instead of the N-terminal serine or cysteine, class 3 inteins contain an additional nucleophilic cysteine residue in Block F. This cysteine in Block F is as part of the unique WCT motif, substituting the function of the N-terminal nucleophilic residue of class 1 inteins required for the first N-S(O) acyl shift step (N-S acyl shift, Fig. 1a)^11,15,16^. Class 3 inteins are thus classified as a distinct class of inteins from class 2 inteins.

**Figure 1.**
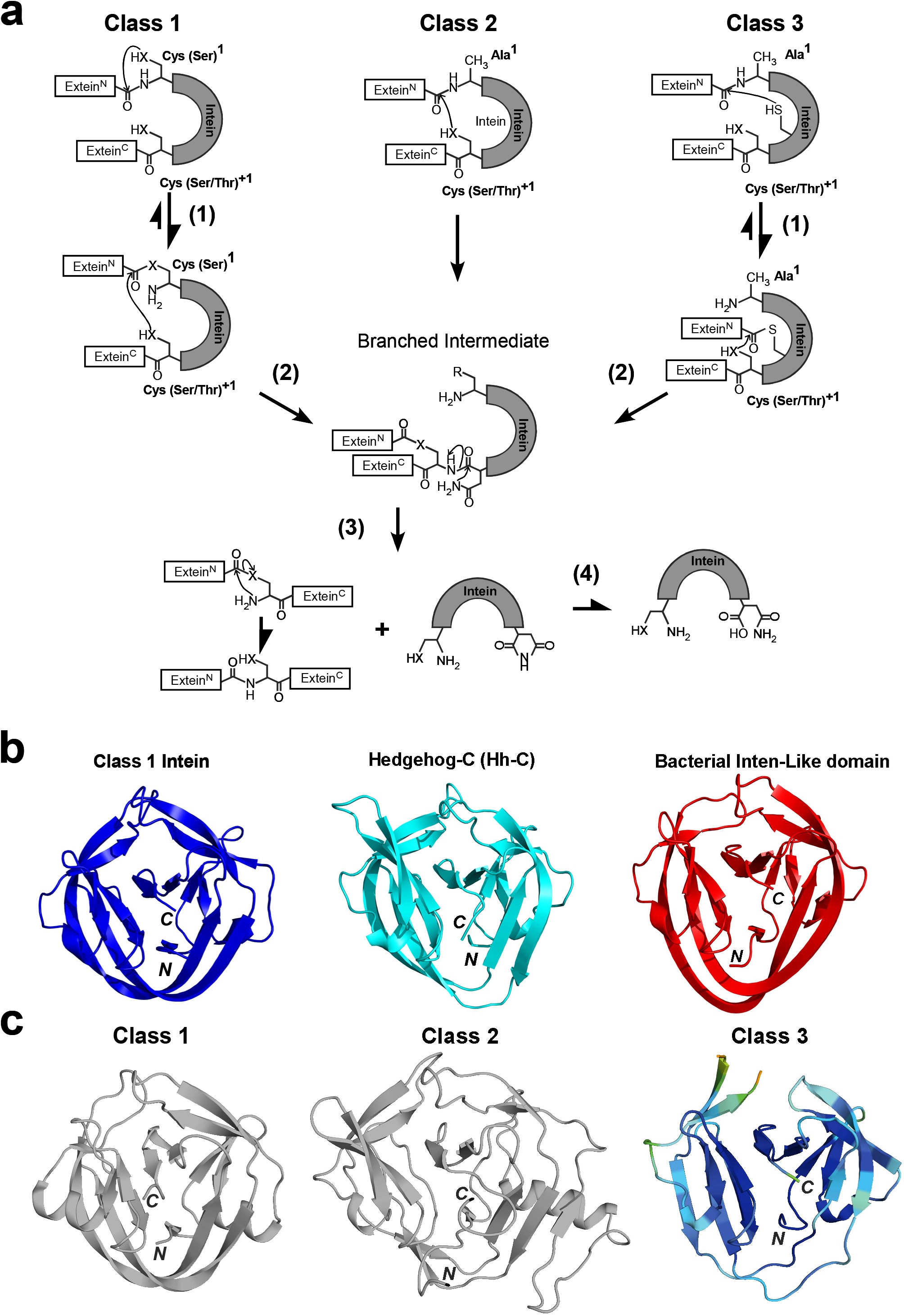
Protein splicing reaction steps for class 1, 2, and 3 inteins^10,11,14,16^. **(a)** Protein splicing mechanisms for class 1 inteins with four concerted steps: (1) N-X acyl shift; (2) *Trans*-(thio)-esterification; (3) Asn cyclization; (4) X-N acyl shift (X = O or S), for class 2 inteins : (2) *Trans*-(thio)-esterification; (3) Asn cyclization; (4) X-N acyl shift (X = O or S), and for class 3 inteins: (1) Thio-ester formation; (2) *Trans*-(thio)esterification; (3) Asn cyclization; (3) N-X acyl shift (X = O or S). **(b)** Ribbon drawing of the structures of representative HINT superfamily members: *Npu*DnaB class 1 intein (4o1r) ^38^, the C-terminal domain of the hedgehog protein (Hh-C, 1at0)^12^, and Bacterial Intein-Like (BIL) domain (2lwy)^27^. **(c)** The crystal structure of class 3 *Mch*Dna1 intein and representative class 1 and 2 inteins. The ribbon drawing of the class 2 intein is based on the *Mja*KlbA intein (2jnq)^58^, and the class 1 intein on the *Npu*DnaE intein (4kl5, chain A)^59^. The ribbon drawing of the *Mch*Dna1 intein (6rix, chain B) structure is colored according to the temperature factor. *N* and *C* denote the N- and C-termini, respectively.

Whereas the first residue for class 1 inteins can be cysteine or serine, the C-terminal nucleophilic residue at the +1 position of inteins is usually either cysteine, serine, or threonine (Fig. 1a). Although the penultimate histidine residue and histidine residue in block B are highly conserved among many inteins, several inteins lack them and remain capable of catalyzing protein splicing by compensatory mutations^19,20^. Inteins catalyzing protein splicing are thus unique single-turnover enzymes that tolerate high sequence variations at the active site residues even among the same class of inteins. Inteins do not have strict requirements for the active site residues but utilize slightly different protein-splicing mechanisms by compensating mutations. Members of the HINT superfamily have been considered to have evolved from a common ancestor by divergent evolution. Although the HINT fold can be easily detected based on the sequence homology, significant deviations of the active-site-residue combinations at all critical residues have been observed^16,18^. How have inteins escaped from the degradation without providing any apparent benefit to their host organisms? How did they evolve into different splicing mechanisms despite the low sequence conservation and high variation of catalytic residues? In this work, we address these questions by revealing the structural basis for the protein splicing mechanism of class 3 inteins by crystal structures, molecular dynamics simulation, and structure-based protein engineering. We propose that the HINT fold could be an effective structural solution for the protein-splicing reaction and an example of protein structure convergence evolved from different ancestral proteins.

## Results

In order to provide an understanding of the class 3 intein splicing mechanism, we decided to determine their structures. We first found that the DnaB1 intein from *Mycobacterium chimaera* (*Mch*DnaB1 intein) has the most robust splicing activity at 37 °C among the tested class 3 inteins from *Deinococcus radiodurans*, *Mycobacterium smegmatis*, and *Mycobacterium chimaera* (Supplemental Fig. S1). We determined the high-resolution crystal structures of two variants of class 3 *Mch*DnaB1 intein (*Mch*DnaB1_HN and *Mch*DnaB1_HAA; Fig. 2 and Supplemental Fig. S2). *Mch*DnaB1_HN (1.66 Å resolution) lacked the C-terminal extein sequence, whereas *Mch*DnaB1_HAA (1.63 Å resolution) contained a C-terminal extein residue (Ala) at the +1 position and a mutation of the terminal Asn residue to Ala (Fig. 1c, Supplemental Fig. S2, Supplemental Table 1). The *Mch*DnaB1 intein structure shares the typical HINT fold of class 1 and class 2 inteins, which is in line with the previous report of class 3 intein structure (Fig. 1b and 1c)^12,23^. Thus, the class 3 *Mch*DnaB1 intein is indistinguishable from class 1 and class 2 inteins by comparing their backbone conformations because additional insertions and deletions observed among inteins easily mask their differences (Fig. 1c)^21^. We found that the most striking feature in the structures of the *Mch*DnaB1 intein is the active site, closely resembling the catalytic triad of serine/cysteine proteases. The observed distance (5.5-5.7 Å) between Sγ and Nδ atoms in the *Mch*DnaB1 inteins is slightly longer than in typical cysteine proteases (3.8-4.0 Å) (Fig. 2a and 2b)^22^. The WCT motif found in the class 3 intein participates in forming the catalytic triad, in which C124, H65, and T143 could serve as nucleophilic, basic, and acidic functional groups, respectively (Fig. 2a and 2b). Importantly, we could observe a large electron density near the side-chains of C124, H65, and the backbone of residue 125 for both crystal structures of the *Mch*DnaDB1_HN and *Mch*DnaB1_HAA inteins (modeled as oxyanion waters in Fig. 2a and Supplemental Fig. S2). This electron density could be the oxyanion hole that is commonly observed in the crystal structures of serine/cysteine proteases, stabilizing the tetrahedral reaction intermediate (Fig. 2a and 2b)^22^. In the class 3 intein structure, Thr143 serves as the protonating acidic residue instead of aspartic acid in the typical Ser-His-Asp catalytic triad of serine proteases. The weaker acidity of Thr compared to Asp might lower not only the nucleophilicity of Cys124 but also increase the distance between His65 and Cys124. However, inteins are single turnover enzymes requiring only one splicing reaction per molecule, rendering high reactivity redundant. Thus, the Cys-His-Thr catalytic triad in *Mch*DnaB1 intein could be sufficient for creating the acyl-enzyme intermediate similar to one found in many serine/cysteine proteases.

**Figure 2.**
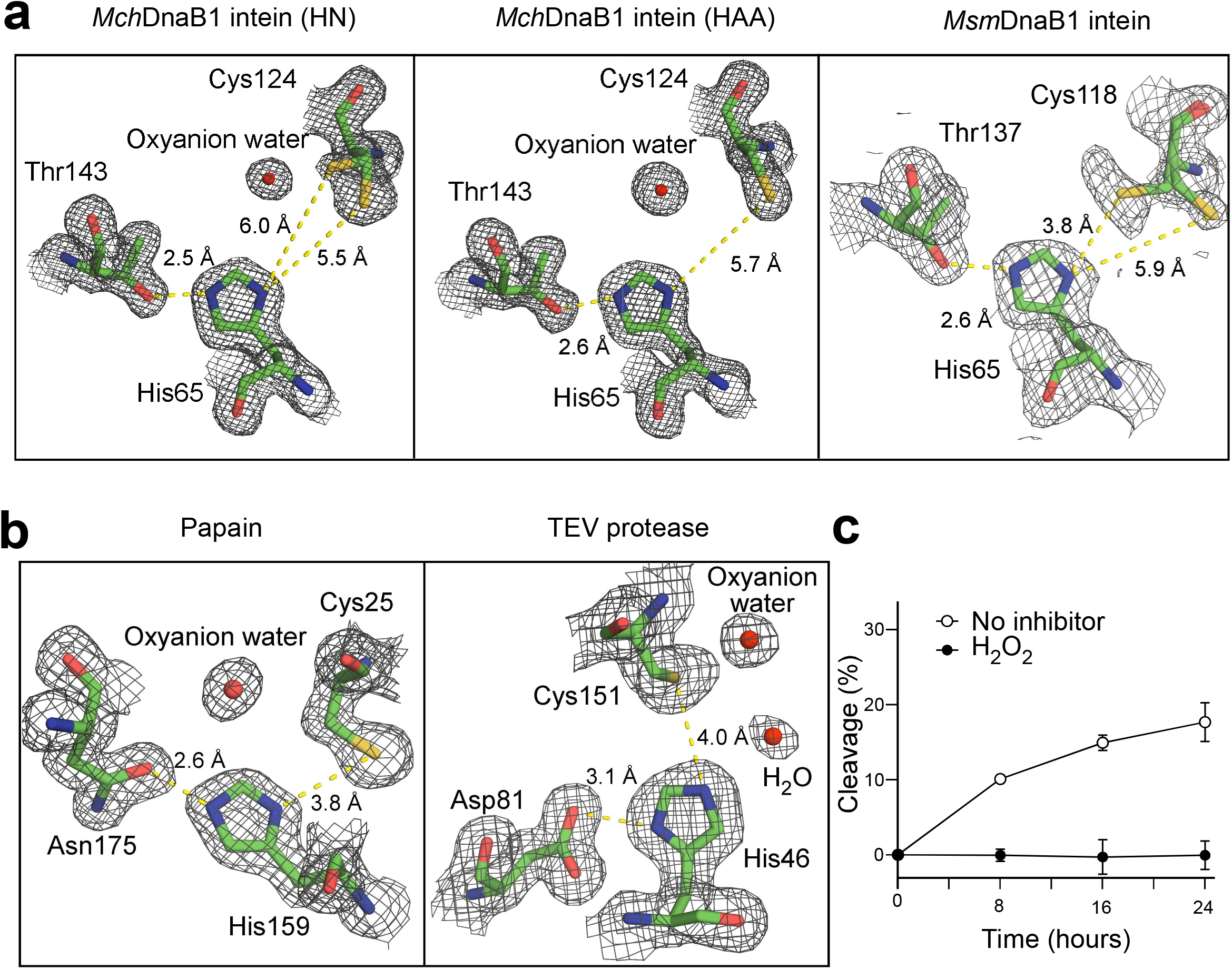
Comparison of the active sites. **(a)** Comparison of the electron density maps at 1.1 sigma counter level around the catalytic-triad between the class 3 inteins: *Mch*DnaB1_HN (6rix), *Mch*DnaB1_HAA (6riy), and *Msm*DnaB1 inteins (6bs8). Oxyanion waters are modeled for the large electron densities near Cys124. **(b)** The electron density maps of the catalytic triad from papain (1ppn) and TEV protease (1lvm). The catalytic triads are depicted together with the electron density maps at the 1.1 sigma counter level. **(c)** Inhibition of the N-cleavage of *Mch*DnaB1_HN by H_2_O_2_. The data were averaged from three replicates. Error bars represent one standard deviation.

### Self-cleavage activity and inhibition of class 3 inteins by protease inhibitors

Both variants of the *Mch*DnaB1 intein were produced for crystallization as N-terminal SUMO fusion proteins, resulting in the N-terminal “SVGK” extein sequence after Ulp1 protease treatment to remove the SUMO fusion tag. However, the crystal structures of both *Mch*DnaB1 intein variants (HN and HAA at the C-terminus) lacked electron densities for the N-extein sequences. This observation is apparently due to self-cleavage at the N terminus during crystallization (N-cleavage)^23^. We also confirmed the N-cleavage activity *in vitro* by incubating the freshly purified fusion proteins over time (Supplemental Fig. S3). As observed for other class 3 inteins, a mutation of the last Asn residue to Ala in the *Mch*DnaB1 intein (*Mch*DnaB1_HAA) largely halted the reaction at the branched acyl-intein intermediate (Supplemental Fig. S3c). Assuming a protease-like mechanism, we tested the inhibition of N-cleavage using common inhibitors of cysteine proteases, phenylmethane sulfonyl fluoride (PMSF) and oxidizing reagent hydrogen peroxide (H_2_O_2_) (Fig. 2c and Supplemental Fig. S3b-c)^24,25^. Whereas PMSF had little effect on the N-cleavage, H_2_O_2_ showed inhibition of the N-cleavage (Fig. 2c and Supplemental Fig. S3b-c). Due to its small size, H_2_O_2_ could easily access to the oxyanion hole, thereby oxidizing Cys124, while PMSF may be sterically-hindered in accessing the active-site cysteine residue, as inteins process an intramolecular substrate. These observations corroborate the notion that a class 3 intein might utilize a catalytic triad similar to serine/cysteine protease for producing the acyl-enzyme intermediate. While most inteins are generally auto-catalytically spliced out immediately after protein translation, the MCM2 intein from *Halorhabdus utahensis* is inactive under low salinity but can be activated at a high salt concentration^26^. To further verify the class 3 splicing mechanism, we used the salt-inducible *Hu*tMCM2 intein for testing the effect of H_2_O_2_ on the N-cleavage of a class 1 intein in an *in vitro* model^26^. We found that H_2_O_2_ did not inhibit N-cleavage of the salt-inducible class 1 intein at a high salt condition, further supporting the protease-like acyl-enzyme intermediate for the class 3 splicing mechanism (Supplemental Fig. S4).

### Conversion of a class 1 intein into a class 3 intein

BIL domains, additional members of the HINT superfamily, predominantly produce N- and C-cleaved products in contrast to protein-splicing domains. BIL domains have probably evolved from inteins divergently^13,27^. We and others previously demonstrated the reverse engineering of BIL domains into efficient *cis*-splicing domains^27,28^. This simple conversion from a BIL domain into a protein-splicing domain implies divergent evolution of BIL domains from an ancestral intein by genetic mutations. Likewise, class 2 inteins lacking Ser or Cys at the N terminus could also efficiently splice after replacement of Ala at the +1 position by Cys or Ser, suggesting a clear evolutionary connection to class 1 inteins^14^.

To examine the divergent evolution of class 3 inteins from class 1 inteins as previously demonstrated with class 2 intein and BIL, we tested the conversion of a class 1 intein into a class 3 intein. Grafting the unique WCT motif found in class 3 inteins to a class 1 intein with the first Cys/Ser to Ala mutation could result in a functional *cis*-splicing intein if they were related by a divergently evolved lineage. We chose the class 1 gp41-1 intein as a model because it already has Thr at the position corresponding to the WCT motif of class 3 inteins and the 1.0 Å-resolution crystal structure is available, facilitating the WCT motif engineering^29^. We grafted the WCT motif to the gp41-1 intein based on the amino-acid sequence alignment (Fig. 3a). However, the engineered class 3 gp41-1 intein (gp41-1_WCT) produced dominantly the C-cleaved product and only a minute amount of the possible splicing product. This result clearly shows that class 3 intein requires additional compensatory mutations in addition to the WCT motif for productive protein splicing (Fig. 3b). To better understand the structural basis for non-productive splicing of the engineered class 3 intein, we solved the crystal structure of gp41-1_WCT at 1.85 Å resolution (Fig. 3c and 3d). Unlike in the crystal structures of *Mch*DnaB1_HAA and *Mch*DnaB1_HN, we observed apparent electron densities for the N-terminal extein, confirming that gp41-1_WCT is inactive in proteolytic cleavage at the N-terminal junction (N-cleavage). The catalytic triad of C124-His65-Thr143 and Trp67 from the WCT motif in the *Mch*DnaB1 intein can be precisely superimposed with the engineered triad of Cys107-His63-Thr123 and with Trp65, except for the χ^1^ angle of the nucleophilic Cys107 (Fig. 3c). The *trans* conformation of Cys107 in gp41-1_WCT is likely to be induced by the presence of the N-extein (see below). Despite successful engineering of the critical WCT motif in the structure of gp41-1_WCT as same as *Mch*DnaB1 intein, gp41-1_WCT failed in productive protein splicing (Fig. 3c). The unsuccessful conversion of a class 3 intein contrasts with the results from the engineering of class 2 intein and BIL domains into class 1-like inteins, in which protein-splicing variants were created by simple mutations. This reverse engineering suggests that a class 3 intein requires additional compensatory mutations in addition to the WCT motif to be proficient in protein splicing. Such simultaneous compensatory mutations on class 1 or 2 inteins together with the WCT motif is an improbable event according to the current survival model of inteins, which are usually inserted near the active site of enzymes essential for host organisms^14,27,28^. A plausible alternative explanation for the emergence of class 3 inteins is that they have gone through a unique evolutionary pathway different from other HINT members.

**Figure 3.**
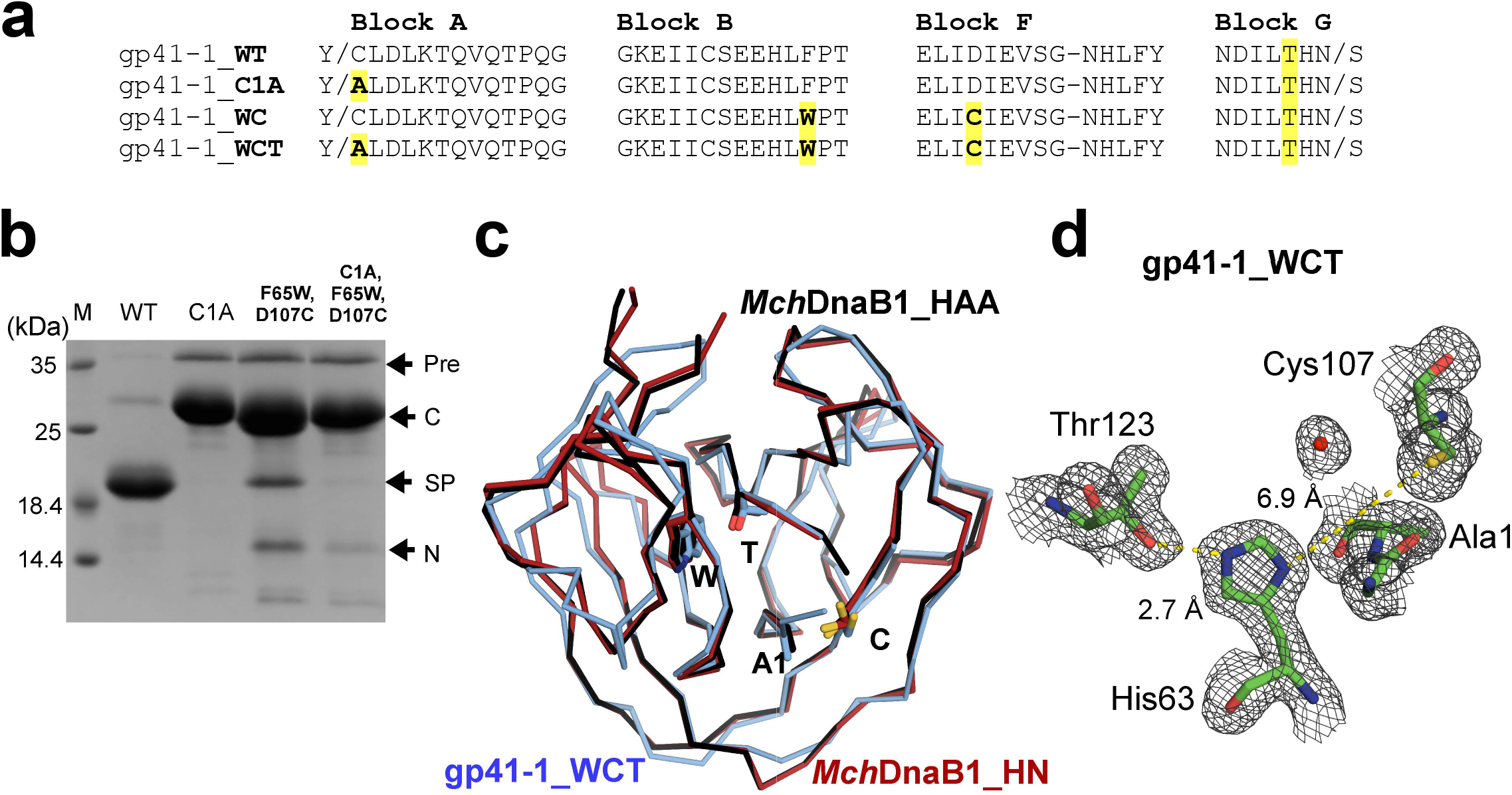
Conversion of a class 1 intein into a class 3 intein by grafting the WCT motif. **(a)** Sequence alignment of the engineered gp41-1 intein variants with different mutations. The WCT motif and C1A substitution are highlighted. **(b)** The SDS-PAGE analysis of protein-splicing by the engineered gp41-1 intein variants with indicated mutations. M, molecular weight marker; WT, wild-type; Pre, precursor; C, C-cleavage product; SP, splicing product; N, N-cleavage product. **(c)** Superposition of the three crystal structures of the gp41-1 intein with the WCT motif (gp41-1_WCT, cyan), *Mch*DnaB1_HN (dark red), and *Mch*DnaB1_HAA (black). The residues of the WCT motif are shown and indicated, together with the first residue of the inteins. **(d)** A close-up of the electron density map observed for the active site of the WCT motif-grafted class 1 intein, gp41-1_WCT.

### The active site of the MchDnaB1 class 3 intein

Despite sharing the same HINT fold, class 3 inteins appear to utilize a very different approach for the same protein splicing reaction in contrast to other members of the HINT superfamily^10,12,16^. Available intein structures containing the extein sequences, except for the two coordinate sets of *Sce*VMA and *Pho*RadA inteins, typically have large distances (~8-9 Å) between the N-scissile peptide and the nucleophilic side chain of the +1 residue that is responsible for the second step, namely *trans*-esterification^30,31,32^. These longer distances suggest the necessity of substantial conformational changes for class 1 inteins during protein splicing. We observed electron density for both the *gauche*+ and *trans-like* conformations of Cys124 in the crystal structure of *Mch*DnaB1_HN, although the side-chain conformation of Cys124 in the *trans-*like conformation is less evident in the second molecule (chain B) in the asymmetric unit (Fig. 2a and Supplemental Fig. S2). A similar conformation was also reported for the structure of another class 3 intein, the DnaB1 intein of *Mycobacterium smegmatis* (*Msm*DnaB1 intein) (Fig. 2a)^23^. On the other hand, the variant of *Mch*DnaB1_HAA shows overall weaker densities for the second conformation in *gauche*+ for Cys124, which was not modeled (Fig. 2a). In the *Mch*DnaB1_HAA intein bearing an extein residue, the distance between the Cβ atom of the +1 residue (Ala) and Sγ atom of Cys124 is 4.7-5.0 Å. However, this distance with the +1 residue of the C-extein would be much shorter (< 3.0 Å) when the χ^1^ angle of Cys124 was in the *trans* conformation. The rotation of the χ^1^ angle of Cys124 could thus bring the nucleophilic atom sufficiently closer to the +1 residue, promoting the *trans*-esterification reaction step without requiring substantial conformational changes reported for other class 1 intein structures^30,31,32^. Therefore, we believe that the movement of Cys124 could play an essential role in the splicing reaction of class 3 inteins, which differs from the reaction mechanisms of class 1 and 2 inteins.

### Molecular Dynamics simulation

To support our interpretation of the *Mch*DnaB1 intein crystal structures, we performed 400-nanosecond MD simulations of *Mch*DnaB1_HN, *Mch*DnaB1_HAA, and the engineered gp41-1_WCT in the presence or absence of the four-residue N-extein. We observed noteworthy differences between the different MD simulations with and without the modeled N-extein for the side-chain conformation of Cys124. The presence of the modeled N-extein pushes the side-chain conformation of Cys124 in both *Mch*DnaB1_HN and *Mch*DnaB1_HAA towards the less favorable *trans*-like conformation (χ^1^ = ~200°-210°) (Fig. 4a, 4b, and Supplemental Fig. S5). Upon removal of the N-extein in the simulation, the population largely shifted towards the ideal *gauche*+ conformation with χ^1^ = ~300° (−60°), with more frequent rotation between *gauche*+ and *trans*-like conformations (Fig. 4b and Supplemental Fig. S5). This observation might suggest that both crystal structures represent the post-splicing or post-cleavage status as expected from the primary structure of the variants (Supplemental Fig. S6). Interestingly, MD simulations also revealed distinct differences between the engineered gp41-1_WCT and *Mch*DnaB1 intein variants. Among the three inteins used for MD simulation, gp41-1_WCT with the N-extein shows the most abundant population for *gauche*- and the χ^1^ angle of the introduced Cys107 is much closer to the ideal 180°*-trans* conformation than to ~200-210° observed in the other simulations for the two *Mch*DnaB1 inteins (Fig. 4a, 4b, and Supplemental Fig. S5). This energetically less favorable *trans*-like conformation observed in *Mch*DnaB1 intein variants might suggest that it could be a driving force for the splicing reaction in class 3 inteins.

**Figure 4.**
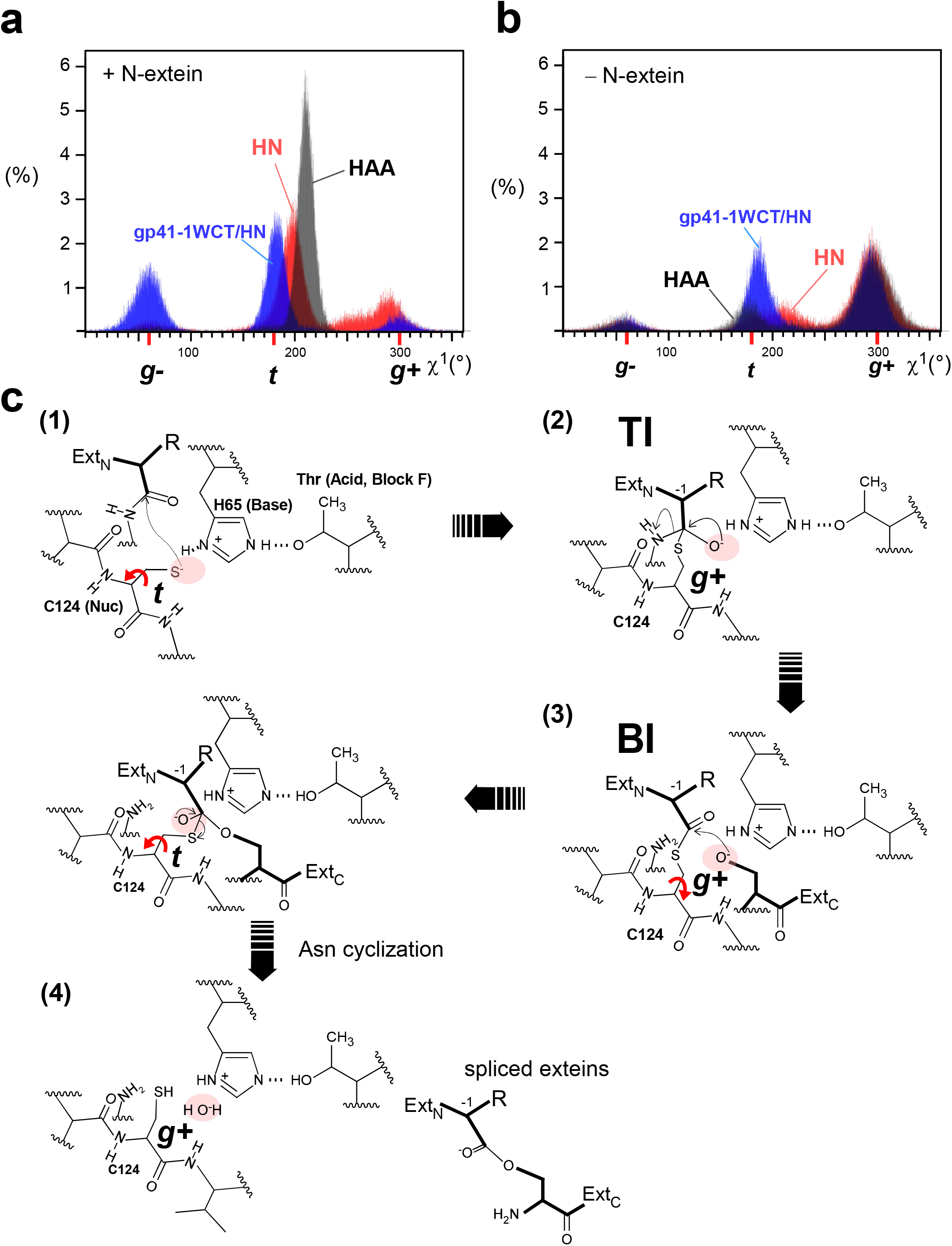
MD simulation and the proposed splicing steps by class 3 inteins. Histograms showing the distributions of the χ^1^ angle for the cysteine residue in the WCT motif during the MD simulations of the two *Mch*DnaB1 intein variants and the engineered gp41-1 intein with WCT motif with the modeled N-extein (**a**) and without N-extein (**b**). Red, grey, and blue colors indicate the data for *Mch*DnaB1_HN, *Mch*DnaB1_HAA, and gp41-1_WCT, respectively. (**c**) Proposed reaction steps for the protein splicing mechanism by the class 3 intein. (1) High energy ground state before splicing. (2) Tetrahedral Intermediate (TI) status after rotation of Cys124 to the gauche^+^ conformation. (3) Branched Intermediate (BI) status. Rotation of Cys124 to the *trans* conformation will bring the thioester intermediate closer to the nucleophilic residue of the C-extein. *Trans*-esterification step via a tetrahedral intermediate. Rotation of Cys124 back to the gauche^+^ conformation. (4) Post-splicing status. Exteins are released from the intein. A red arrow indicates a rotational movement of Cys124.

### The catalytic mechanism of class 3 inteins

Based on biochemical and structural data as well as MD simulations, we propose the catalytic mechanism of class 3 inteins, as depicted in Fig. 4c. At the pre-splicing state, Cys124 is at the high-energy (unfavorable) *trans*-like conformation and is weakly deprotonated by His65. The rotation around the χ^1^ angle of Cys124 to *gauche*+ from the high-energy state would induce the first step of the nucleophilic attack and form the tetrahedral intermediate (TI), which is supposedly stabilized by the oxyanion hole. The subsequent N-cleavage creates a thioester bond in the branched intermediate (BI). The rotation of the χ^1^ angle would bring the branched intermediate bearing the thioester bond closer to the nucleophilic oxygen atom of Ser at the +1 position for the *trans*-esterification reaction *via* the tetrahedral intermediate that might also be stabilized by the oxyanion hole. The more frequent χ^1^ rotation of Cys124 between the *gauche*+ and *trans*-like conformation in the absence of the N-extein could thus mimic the movement of the branched intermediate bringing the state closer to the +1 residue. The subsequent *trans*-esterification reaction via the tetrahedral intermediate stabilized by the oxyanion hole releases the N- and C-exteins from the intein. The released extein ester will undergo subsequent O-N rearrangement to the energetically favorable peptide bond. Based on our current data, it is unclear whether Asn cyclization will take place prior to the *trans*-esterification or simultaneously with it. The intein reaches the ground state of the *gauche*+ of Cys124, represented by the crystal structure of *Mch*DnaB1_HN. In the absence of the nucleophilic +1 Ser residue, the oxyanion water molecule slowly hydrolyzes the branched intermediate and releases the N-extein. We think that the three-dimensional crystal structure of *Mch*DnaB1_HAA likely represents the post-hydrolysis state of the *Mch*DnaB1 intein (Supplemental Fig. S6). In this proposed model for the splicing mechanism of the class 3 intein, the rotational motion of the cysteine in the WCT motif might play a critical role, unlike in other intein classes where wide conformational changes of 8-9 Å are expected to occur for the first N-S(O) acyl shift^30,32^.

## Discussion

One protein fold may serve as a common scaffold for many functions. For example, the eightfold (βα) barrel structure, known as TIM-barrel, is the most common protein fold utilized by many different enzymes with very diverse amino-acid sequences^33^. Whereas a specific protein fold might not be a prerequisite for the function of a protein, the catalytic triad found in proteases is often considered as a prime example of convergent evolution^2^. This convergent evolution is assumed because it is unlikely that two proteins evolving from a common ancestor could have retained similar active-site structures while other structural features have completely changed^1^. Many serine/cysteine proteases, such as chymotrypsin/trypsin, share the two-barrel motif as the core – a result of presumable gene duplication (Fig. 5)^34^. The acid-histidine-nucleophile catalytic triad motif of serine/cysteine proteases is located at the interface of the two β-barrels and considered to be the result of convergent evolution^3^. Even though the common horseshoe-like fold of the HINT superfamily members, including inteins, does not have two distinct β-barrels, the HINT fold contains two subdomains related by the pseudo-C2-related symmetry^12^. This symmetry relation is considered to be the result of gene duplication, fusion, and loop-swapping events^12,34^. The catalytic triad formed by Cys124-His64-Thr143 in class 3 *Mch*DnaB1 intein is analogously split between the two subdomains and located at the interface. The catalytic triad being at the interface of the two subdomains of the HINT fold resembles the common catalytic triad of serine/cysteine proteases, including the oxyanion hole stabilizing the tetrahedral intermediate during catalysis (Fig. 2a). Since peptide bond formation is the reverse reaction of peptide hydrolysis, it is not surprising that protein splicing uses the same mechanism as cysteine proteases involving a tetrahedral intermediate. Indeed, several peptidases have been used for *trans*-peptidase reactions^35,36^.

**Figure 5.**
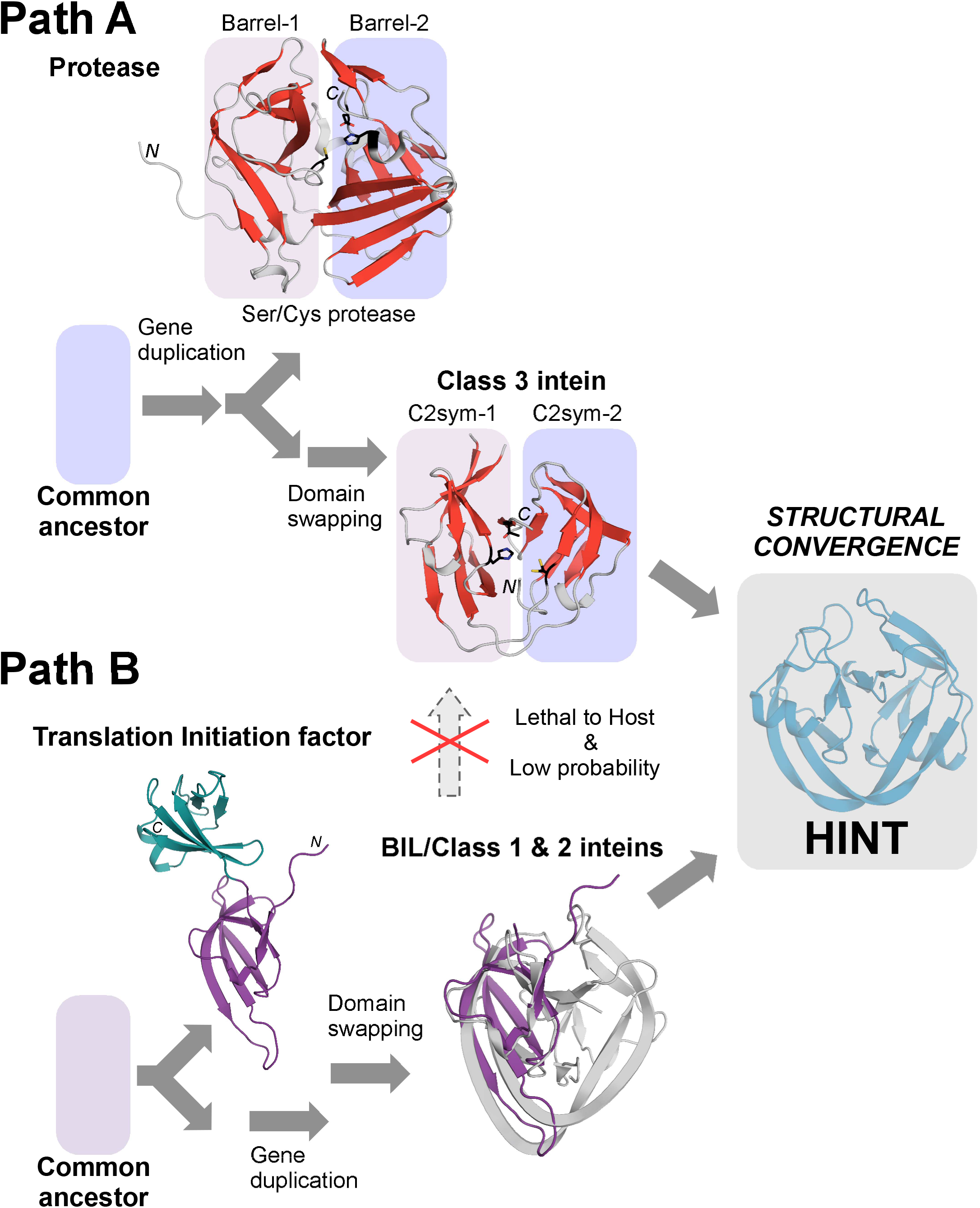
Convergent evolution model of the HINT fold. (**a**) Class 3 inteins may have evolved from an ancestral cysteine protease originating from prophages, retaining the highly conserved catalytic triad. Two domains of a cysteine protease (TEV protease, 1lvm) and the pseudo-C2-symmetry relation in a class 3 intein (*Mch*DnaB1 intein, 6rix) are indicated with circles. (**b**) Other members of the HINT superfamily might have evolved via a very different pathway from a distantly related ancestral protein such as a translation initiation factor by gene duplication, fusion, and domain swapping. Cartoon drawings of translation initiation factor (IF5A, 1bkb)^39^ and the superposition with the pseudo-C2-related subdomain of the BIL domain (2lwy)^27^ are shown (**Supplemental Table S2**). The purple N-terminal domain of IF5A was superimposed with the HINT domain.

A comparison between the splicing active *Mch*DnaB1 intein and the WCT motif-engineered inactive gp41-1 intein derived from a class 1 intein strongly implies that accumulation of random mutations in a class 1 intein would not directly lead to a class 3 intein. Such divergent evolution model of class 3 inteins is particularly implausible because any functionally detrimental mutations of the active site residues of inteins could reduce the fitness of the host organism or even be lethal. Concurrent compensatory mutations constantly maintaining the splicing activity is an improbable event, suggesting that class 3 cannot be directly evolved from a class 1 or 2 intein.

The MD simulations provided additional evidence that the rotational motion of the active-site cysteine could be sufficient for enabling protein splicing of class 3 inteins, unlike class 1 and 2 inteins, which require substantial conformational changes. Class 3 inteins hence utilize a different catalytic mechanism. The WCT motif engineering on a class 1 intein does not seem to provide similar rotational dynamics of the active site residue, indicating that additional compensatory mutations are required for splicing active inteins. The structural and biochemical results impose the question of how class 3 inteins could have divergently emerged from class 1 or class 2 inteins. A plausible explanation from the structural basis of class 3 splicing mechanism is that class 3 inteins have evolved from a protease-linage originating from prophages different from other class 1 and 2 inteins^8,11,16,19,20^.

Inteins tolerate a vast array of variations at the active site for protein splicing, leaving the N-terminal Ser/Cys and C-terminal Asn/Gln/Asp as the only omnipresent amino-acid residues among class 1 inteins because even a highly conserved histidine in block B and penultimate histidine are substituted in several inteins^19,20^. These conserved residues can be further reduced to the C-terminal Asn for class 2 inteins, yet retaining the protein splicing activity by different combinations of the catalytic residues and compensatory mutations. One way to explain the extremely high tolerance of the active sites of inteins is that the HINT fold is the critical structural solution enabling peptidyl transfer reactions. In the HINT fold, the enzymes (inteins) and substrates (exteins) are covalently connected as single precursor molecules, thereby working as single-turnover enzymes. Inteins do not involve any substrate-association step. The covalent linkage to its substrates could also facilitate the accommodation of different amino-acid types at the active site residues among the HINT superfamily compared with other enzymes. The critical role of the HINT fold is to bring the acyl-(thio)ester intermediate and the nucleophilic residue from the C-extein close together, at the precise position and timing required for protein splicing. We gathered evidence suggesting that class 3 inteins might have evolved through a different pathway than class 1 and 2 inteins, possibly related to serine/cysteine proteases originated from prophage because class 3 inteins have a clear monophyletic distribution and inactive class 3 intein sequence was found within a pseudogene^16,17^. We revisited what would be the possible common ancestral protein of other members among the HINT superfamily. We searched with the BIL coordinates (2lwy)^27^ the Protein Data Bank (PDB) using DALI server^37^ and identified possible ancestral domains corresponding to the C2-related pseudo-symmetry subdomain in the HINT fold (Supplemental Table S2). Despite their low Z-scores (2.5-2.7), we noticed structural homology to translation initiation factor 5A (1bkb)^39^, eukaryotic translation initiation factor 5A2 (3hks)^40^, and elongation factor P (1ueb)^41^, demonstrating the apparent structural similarity with r.m.s.d. values between 1.8 and 2.4 Å for 42-49 residues (Fig. 5 and Supplemental Figure S7). Intriguingly, these proteins are also involved in the first step of peptide bond formation in translation utilizing ribosomal protein synthesis. Class 1 and 2 inteins might have descended from a common ancestor shared by translation initiation factors or their ancestor by gene duplication and swapping^12^.

In summary, we identified possible convergent evolution of the HINT fold in protein splicing by deconvoluting the existing intein structures and their reaction mechanism into possible ancestral proteins with distantly related origins. Despite the identical HINT fold, the protein-splicing mechanisms seem to have widely diverged, which cannot be explained by the divergent evolution model by random mutations, because inteins would require several concurrent compensatory mutations for their survival. The extremely high diversity of the active-site residue combinations found in protein splicing could be reminiscent of independent evolutionary pathways originating from the distantly related ancestral proteins shared with proteases and translation initiation factors, yet leading to the same structural solution, i.e., the HINT fold. We propose that the HINT fold is an effective structural and biochemical solution for *trans*-peptidyl reactions and the first example structural convergence of a whole protein. Deconvoluting functional mechanisms and ancestral structural protein domains might assist in identifying further examples of structural convergence of various other protein folds.

## Methods

### Cloning of class 3 intein expression vectors

The gene encoding the *Mch*DnaB1 intein () was amplified from the genomic DNA of *Mycobacterium chimaera* strain DSM 44623 using two the oligonucleotides HB095: 5’-GTGGATCCGTCGGGAAGGCCCTTGC and HB096: 5’-CTGGGTACCTAGCGTGGAATTGTGCGTCG. The amplified gene was cloned between the *Bam*HI and *Kpn*I sites of pSKDuet16^42^, resulting in pHBDuet071 for *cis*-splicing tests. The gene was further PCR-amplified from pHBDuet071 using the two oligonucleotides J765: 5’-GAACAGATTGGTGGATCCGTCGGGAAGGCCCTTGC and J759: 5’-GTGCGGCCGCAAGCTTAATTGTGCGTCGGCACCATCCCGC for *Mch*DnaB1_HN, or J765 and J760: 5’-GTGCGGCCGCAAGCTTAGGCAGCGTGCGTCGGCACCATCCCGCG for *Mch*DnaB1_HAA. The PCR products were ligated into *Bam*HI and *Hin*dIII-digested pHYRSF53^43^, resulting in pHBRSF073 (*Mch*DnaB1_HN) and pHBRSF074 (*Mch*DnaB1_HAA) for the bacterial expression of N-terminally hexahistidine-tagged SUMO-fused *Mch*DnaB1 intein variants.

The C1A, F65W, and D107C mutations were introduced into the gp41-1 intein coding sequence via assembly PCR from plasmid pBHDuet37^29^ using the oligonucleotides HB019: 5’-CAAAACCTACACCGTAACGGAAGGATCCGGCTATGCGCTGGATCTGAAAACGCAGGTGC and HB015: 5’-CGGTCTGGGTCGGCCACAGATGTTCTTCGCTACAAATAATTTCTTTG, HB016: 5’-CGAAGAACATCTGTGGCCGACCCAGACCGGCGAAATG and HB017: 5’-CGCTCACTTCAATGCAGATCAGTTCGCGTTCATCCAGCTC, HB018: 5’-GAACGCGAACTGATCTGCATTGAAGTGAGCGGTAACCATCTG and HB014: 5’-CGTTCAGGATAAGTTTGTACTGGGTACCGCTCGAGCTGTTGTGGGTCAGAATGTCGTTC, thereby attaching the 3-residue N- and C-terminal junction sequences. The assembled PCR product was ligated into pBHDuet37^29^ using the *Bam*HI and *Kpn*I restriction sites, resulting in plasmid pHBDuet024 for *cis*-splicing tests. Variants encoding only the F65W and D107C mutations (pHBDuet023) were generated the same way, but using HB013: 5’-CAAAACCTACACCGTAACGGAAGGATCCGGCTATTGCCTGGATCTGAAAACGCA GGTG instead of HB019. For introducing the C1A mutation (pHBDuet022), the oligonucleotides HB019 and HB014 were used. As gp41-1 *cis*-splicing wild-type control, plasmid pHBDuet021^29^ was used. For structural studies on gp41-1_WCT, the gene was amplified from pHBDuet024 using the oligonucleotides I521: 5’-TTGGATCCGGTGGTGCCCTGGATCTGAAAACGCAG and I522: 5’-GTCAAGCTTAGTTGTGGGTCAGAATGTCGTTC and ligated into *Bam*HI and *Hin*dIII-digested pHYRSF53^43^ resulting in pHBRSF044 encoding N-terminally hexahistidine-tagged SUMO-fused gp41-1_WCT. Plasmid pET22b_TRX_MSM encoding the *Msm*DnaB1 intein was a kind gift from Dr. FB. Perler (New England Biolabs, USA). The intein gene was amplified using the oligonucleotides HK960: 5’-AGGGATCCGGTAAAGCACTGGCACTGGAT and HK961: 5’-AGCAAGCTTAGGTCGCATTATGGGTCGGAACCATACC and ligated into *Bam*HI and *Hin*dIII-digested pHYRSF53^43^ resulting in pCARSF64, encoding N-terminally hexahistidine-tagged SUMO-fused *Msm*DnaB1_HNAT intein with three N-terminal Ser-Gly-Lys, and two C-terminal Ala-Thr extein residues. Alternatively, HK960 was used with HK971: 5’-AGCAAGCTTAATTATGGGTCGGAACCATACC, resulting in pCARSF63-65 lacking the C-terminal extein sequence. pCARSF63-65 was used for the production of ^15^N-labeled *Msm*DnaB1. The *cis*-splicing vector pHBDuet060 was constructed by PCR amplification of the *Msm*DnaB1 intein from pCARSF64 using the oligonucleotides HB078: 5’-GGAAGGATCCGTGGGTAAGGCGCTCGCGCTCGACAC and HB079: 5’-ACTGGGTACCGAGTGTCGAGTTGTGCGTGGGAACCATG and ligation of the product into pSKDuet16^42^ using *Bam*HI and *Kpn*I sites.

The gene encoding the *Dra*Snf2 intein was amplified from the genomic *Deinococcus radiodurans* DNA (DSM-20539) using the oligonucleotides HB020: 5’-GAAGGATCCCTGGGCAAGGCGCAGC and HB021: 5’-ACTGGGTACCTTGCAGCGTGTTGTGGGTG including three residues of N- and C-terminal junction sequence. The PCR product was ligated into pSKDuet16^42^ using *Bam*HI and *Kpn*I sites, resulting in the *cis*-splicing vector pHBDuet027. The nested endonuclease domain was deleted by PCR amplification of the N- and C-terminal halves using HB020 and HB072: 5’-CGCTGCCGCCGCTGCCACTGCCACCGCTGCCACTACCGCCGGGGTCGAGGGGCAG, and HB071: 5’-CGGTGGCAGTGGCAGCGGCGGCAGCGGTGGCAGTGGCAGCGGCGGCGAGAAGAAAACG and SZ015: 5’-TGCCAAGCTTATTCCGTTACGGTG and assembled with HB020 and SZ015. The product was ligated into pSKDuet16^42^ as described above, resulting in the *cis*-splicing vector pHBDuet058 encoding the *Dra*Snf2 intein with a deletion of residues 121-266 replaced by an 18-residue GS-based linker (*Dra*Snf2^Δ128^). For deleting residues 121-251 (*Dra*Snf2^Δ131^, pHBDuet057), the oligonucleotides HB069: 5’-GCGGGCCACCCCGCCGGGGTCGAGGGGCAG, and HB070: 5’-CTGCCCCTCGACCCCGGCGGGGTGGCCCGCATTC were used instead of HB072 and HB071. For testing salt-inducible N-cleavage of a class 1 intein, plasmid pSADuet735 was used encoding the *Hut*MCM2 intein with the terminal and +1 intein residues mutated to Ala, flanked by two GB1 domains and N-terminal hexahistidine tag (H_6_-GB1-*Hut*MCM2_HAA-GB1)^26^. All the plasmids used, except for pSADuet735, are deposited at www.addgene.org (www.addgene.org/Hideo_Iwai).

### Production and purification of *Mch*DnaB1_HN, *Mch*DnaB1_HAA, and gp41-1_WCT

Proteins were produced in *E. coli* strain T7 Express (New England Biolabs, Ipswich, USA) in 2 L 25 μg mL^−1^ kanamycin-containing LB medium by induction with 1 mM isopropyl-β-D-thiogalactoside (IPTG). *Mch*DnaB1_HAA and gp411_WCT were expressed at 37 °C for 3 hours, *Mch*DnaB1_HN at 16 °C overnight. The induced cells were harvested by centrifugation at 4700×*g* for 10 min at 4 °C and frozen in liquid nitrogen for storage at −80°C. The harvested cells were lysed in buffer A (50 mM sodium phosphate pH 8.0, 300 mM NaCl) using continuous passaging through an EmulsiFlex-C3 homogenizer (Avestin, Mannheim, Germany) at 15,000 psi for 10 min, 4 °C. Lysates were cleared by centrifugation at 38000 ×*g* for 60 min, 4 °C. Proteins were purified in two steps by immobilized metal chelate affinity chromatography (IMAC) using 5 mL HisTrap FF columns (GE Healthcare, Chicago, Illinois, USA) as previously described, including the removal of the hexahistidine tag and SUMO fusion^43^. During the two IMAC purification steps, proteins were dialyzed against the following buffers: *Mch*DnaB1_HN, buffer B (phosphate buffer saline (PBS) supplemented with 100 mM NaCl, 1 mM dithiothreitol (DTT)) and Buffer C (20 mM Tris-HCl pH 8.0, 200 mM NaCl, 1 mM DTT); *Mch*DnaB1_HAA, PBS and Buffer D (10 mM Tris-HCl pH 7.5, 100 mM NaCl, 1 mM DTT); gp41-1_WCT, PBS and deionized water. *Mch*DnaB1_HN and gp41-1_WCT were further purified using a Superdex® 75 10/300 column (GE Healthcare, Chicago, Illinois, USA) in buffer E (10 mM Tris-HCl pH 8.0, 200 mM NaCl, 1 mM DTT) and Buffer F (0.5× PBS, 1 mM DTT), respectively. Peak fractions were combined, dialyzed, and concentrated using Macrosep® Advance Centrifugal Devices 3K MWCO (Pall, Port Washington, USA), and used for crystallization trials.

### Proteolytic inhibition assays

The class 3 intein constructs H_6_-SUMO-*Mch*DnaB_HN and H_6_-SUMO-*Mch*DnaB1_HAA, and the salt-inducible class 1 intein H_6_-GB1-*Hut*MCM2_HAA-GB1 were produced in *E. coli* strain T7 Express (New England Biolabs, Ipswich, USA) at 37 °C in 5 mL LB medium containing 25 μg mL^−1^ kanamycin by induction with 1 mM IPTG for 3 hours. The induced cells were harvested by centrifugation at 4700 ×*g* for 10 min, and proteins were purified by IMAC using Ni-NTA spin columns (QIAGEN, Hilden, Germany). Proteins were eluted in 100 μL elution buffer (50 mM sodium phosphate pH 8.0, 300 mM NaCl, 250 mM imidazole) and incubated after addition of 1 or 10 mM phenylmethanesulfonyl fluoride (PMSF) (Roche, Basel, Switzerland), or 1 mM H_2_O_2_ (Sigma Aldrich, Steinheim, Germany) at room temperature (RT) for class 3 inteins. The N-cleavable salt-inducible class 1 intein was incubated in 0.35 M sodium phosphate buffer pH 7.0, 3.5 M NaCl, 0.5 mM EDTA at RT. Samples were taken at the indicated time points and analyzed by SDS-PAGE (16.5%). Band intensities were quantified using ImageJ 2.0.0-rc-69/1.52p. The quantification of the N-cleavage of H_6_-SUMO-*Mch*DnaB1_HN and H_6_-GB1-*Hut*MCM2_HAA-GB1 was derived from the equation, 100 × [(CP/(CP+P))_t_ - (CP/(CP+P))_t0_]/[1 - (CP/(CP+P))_t0_], where CP is the sum of the cleavage products (H_6_-SUMO and *Mch*DnaB1_HN, or H_6_-GB1 and *Hut*MCM2_HAA-GB1), t and t0 are the time and zero-time points, and P is the unreacted precursor.

### Protein cis-splicing tests

To assay protein *cis*-splicing, the vectors encoding the inteins *Msm*DnaB1 (pHBDuet060), *Mch*DnaB1 (pHBDuet071), *Dra*Snf2 (pHBDuet027), *Dra*Snf2^Δ128^ (pHBDuet058), and *Dra*Snf2^Δ131^ (pHBDuet057) were expressed in *E. coli* strain T7 Express (New England Biolabs, Ipswich, USA) as described in section “Proteolytic inhibition assays”. Proteins were purified and analyzed as described above.

### Crystallization and structure determination of MchDnaB1_HN, MchDnaB1_HAA, gp41-1_WCT

Diffracting crystals of *Mch*DnaB1_HN were obtained at room temperature by mixing 100 nL concentrated protein (13.4 mg/mL) with 100 nL mother liquid (100 mM Tris-HCl pH 9, 200 mM MgCl_2_, 30% (w/v) polyethylene glycol (PEG) 4000). Data were collected at beamline i03 at Diamond Light Source (Didcot, UK) equipped with a Pilatus detector. Data were processed to 1.66 Å (Supplemental Table 1). The structure was solved by molecular replacement using PHASER with the *Msm*DnaB1 intein (6bs8) as a search model^44,23^. The model was built using PHENIX AutoBuild, manually corrected with COOT, and refined using PHENIX^46^. The final model consists of two molecules in the asymmetric unit. The four residues of the sequence SVGK preceding Ala1 of the intein were clearly missing in the electron density. A loop region between residues Ser91 - Leu104 (chain A) and Gly90 - Leu104 (chain B) was not modeled due to insufficient density information. The electron density for the side chain Cys124 suggested that it was oxidized and was therefore modeled as S-oxy cysteine (Csx). Alternate conformations were modeled for Thr15, Asp19, Arg46, and Csx124 (chain A) and Cys124 and His144 (chain B). The final model includes one Cl^−^ ion originating from the crystallization buffer. The structure was validated using MolProbity (score 1.07, 100^th^ percentile)^47^.

*Mch*DnaB1_HAA crystals were obtained as described above using concentrated protein (13 mg/mL) after adjusting the DTT concentration to 10 mM and mother liquid (100 mM Tris-HCl pH 7.5, 200 mM MgCl_2_, 25% (w/v) PEG 4000). Data were collected at beamline ID30A-1 / MASSIF-1 at ESRF (Grenoble France) equipped with a Pilatus detector and processed to 1.63 Å. (Supplemental Table S1). The structure was solved by molecular replacement using PHASER with the *Mch*DnaB1_HN structure as a search model^44^. The structure model was built using ARP/wARP, manually corrected using COOT and refined using PHENIX^45,46, 49^. The final model consists of two molecules in the asymmetric unit. Four residues of the sequence SVGK preceding Ala1 of the intein were clearly missing in the electron density. A loop region between residues Gly90 - Leu105 (chain A) and Gly90 - Leu104 (chain B) was not modeled due to the lack of electron densities. Alternate conformations were modeled for Thr15, Pro142, (chain A), and Val87 (chain B). The final model contains one Cl^−^ ion. The structure was validated using MolProbity (score 1.04, 100^th^ percentile)^47^.

Diffracting crystals of gp41-1_WCT were obtained as above with a protein concentration of 40 mg/mL and mother liquid (100 mM bis-tris pH 5.5, 200 mM (NH_4_)_2_SO_4_), 25% (w/v) PEG 3350). Data were collected at beamline I04 at Diamond Light Source (Didcot, UK) equipped with a Pilatus detector and 1.85 Å (Supplemental Table S1). The structure was solved by molecular replacement using PHASER with the gp41-1 intein (6qaz) as a search model^44^. The structure model was built using PHENIX AutoBuild, manually corrected with COOT and refined using PHENIX^46,45^. The entire protein chain (one molecule in the asymmetric unit) could be traced in the electron density without breaks for all 128 residues except for the first Ser residue. A non-canonical *cis* peptide bond was modeled between Lys87 and Glu88, which is also found in the search model. Alternate conformations were modeled for Leu25, Ser28, Val38, and Ser46. Additional density was observed for the sidechain of Cys83 indicating oxidation and was modeled as 3-sulfinoalanine (Csd). The structure was validated using MolProbity (score 1.28, 99^th^ percentile) ^47^.

### Molecular Dynamics Simulation

We performed MD simulations of the three different proteins, *Mch*DnaB1_HN, *Mch*DnaB1_HAA, and gp41-1_WCT, with and without modeling an N-extein. The crystal structures of both *Mch*DnaB1_HN (chain B) and *Mch*DnaB1_HAA (chain A) were missing residues in a loop region (see above). After modeling the missing residues with MODELLER software^49^, the crystal structures were used as the starting structure for the simulation without the N-terminal residues. The four-residue N-extein (“SVGK”) was modeled on the structure to generate the initial structure for the MD simulation with the N-terminal residues using the MODELLER software^49^. The crystal structure of gp41-1_WCT (6riz) contained all the residues, including the N-extein part of the “GG” sequence, and it was used as the starting structure for the simulation with the N-extein part. The initial structure of the gp41-1_WCT simulation without the N-extein part was derived by removing the first two glycine residues from the crystal structure.

The MD simulations were performed using Gromacs 2018 software^51^ and Amber ff99SB-ILDN force field^52^ in a rectangular simulation box with periodic boundary conditions. The protein coordinates from the crystal structures of *Mch*DnaB1_HN, *Mch*DnaB1_HAA, and gp41-1_WCT were solvated with approximately 11,000 and 7,500 TIP3P water molecules^53^, and the systems were made electroneutral by adding an appropriate number of Na^+^ ions. The structures were first energy minimized for 1000 steps with the steepest descent algorithm. The production simulations were run for 400 ns with a timestep of 2 fs for each system. All bond lengths were constrained with LINCS^54^. The temperature was set to 303K with the v-rescale thermostat^55^, and Parrinello–Rahman barostat was used for isotropic pressure coupling at 1 bar^56^.

Electrostatic interactions were treated with particle mesh Ewald^56,57^, and Lennard-Jones interaction cut-off was set to 1.0 nm. The χ1 angle of the cysteine residue within the active site (Cys124 for *Mch*DnaB1_HN and *Mch*DnaB1_HAA, and Cys107 for gp41-1_WCT, respectively) was analyzed with Gromacs utilities. The simulation data are available from the Zenodo repository (DOI:10.5281/zenodo.3448608).

## Abbreviations

CBD: chitin-binding domain
BI: branched intermediate
BIL: Bacterial Intein-Like
HINT: Hedgehog/INTein
Hh-C: the C-terminal domain of the Hedgehog protein or hog protein
IMAC: immobilized metal affinity chromatography
IPTG: isopropyl-β-D-thiogalactoside
*Mch*DnaB1 intein: DnaB1 intein from *Mycobacterium chimaera*
PDB: Protein Data Bank
r.m.s.d.: root-mean-square deviation
PEG: polyethylene glycol
PMSF: phenylmethane sulfonyl fluoride
DTT: dithiothreitol

## Acknowledgments

This work is supported in part by the Academy of Finland (1277335, 315596), Novo Nordisk Foundation (NNF17OC0025402, NNF17OC0027550), and Sigrid Jusélius Foundation, as well as by the Intramural Research Program of the NIH, National Cancer Institute, Center for Cancer Research and with Federal funds from the National Cancer Institute, National Institutes of Health, under Contract No. HHSN261200800001E (to GTL). The Finnish Biological NMR Center is supported by Biocenter Finland and HiLIFE-INFRA. We acknowledge CSC–IT Center for Science (Finland) for computational resources. We thank T. V. Kudling and Dr. V. Manole for their assistance in protein production and crystallization. The content of this publication is solely the responsibility of the authors and does not necessarily represent the official views or policies of the Department of Health and Human Services, nor does the mention of trade names, commercial products, or organizations imply endorsement by the U. S. Government.

