## Supplemental Table 1 for "Convergent evolution of the Hedgehog/Intein fold in protein splicing"

**Supplemental Table 1** Data collection and refinement statistics.

| | | | |
|---|---|---|---|
| **Data collection** | | | |
| Intein | *Mch*DnaB1_HN | *Mch*DnaB1_HAA | gp41-1_WCT |
| Beamline | i03 (Diamond) | ID30A-1 (ESRF) | i04 (Diamond) |
| Program | HKL3000 | Mosflm | HKL3000 |
| Wavelength | 0.9763 | 0.966 | 0.9795 |
| Space group | $P2_1$ | $P2_1$ | $P6_522$ |
| Molecules / a.u. | 2 | 2 | 1 |
| Unit cell *a*, *b*, *c* (Å); | 31.25, 95.34, 35.79 | 31.31, 95.78, 35.86 | 90.22, 90.22, 71.73 |
| $\alpha$, $\beta$, $\gamma$ (°) | 90, 107.14, 90 | 90, 107.37, 90 | 90, 90, 120 |
| Resolution range (Å) | 50.0 - 1.66 (1.68 - 1.66) | 34.23 - 1.66 (1.66-1.63) | 50 - 1.85 (1.88 - 1.85) |
| Total no. of reflections | 135528 | 60056 | 509713 |
| No. of unique reflections | 22641 (731) | 24075 (1195) | 15226 (741) |
| $R_{merge}$ (%)$^{\dagger}$ | 8.5 (36.7) | 8.1 (50.1) | 11.7 (77.3) |
| $<I/\sigma_I>$ | 18.7 (2.2) | 7.3 (1.9) | 33 (2.7) |
| $CC_{1/2}^{\&}$ | 0.993 (0.543) | 0.994 (0.603) | 1.000 (0.554) |
| Completeness (%) | 95.0 (60.2) | 95.8 (96.8) | 100 (99.1) |
| Redundancy | 6.0 (2.7) | 2.5 (2.6) | 33.5 (14.3) |
| **Refinement** | | | |
| Resolution range (Å) | 47.67 - 1.66 (1.74 - 1.66) | 34.23- 1.63 (1.70 - 1.63) | 45.11 - 1.85 (1.99 - 1.85) |
| No. of reflections (refinement / $R_{free}$) | 21521 / 1037 | 24035 / 1186 | 15147 / 717 |
| $R$ / $R_{free}$ (%)$^{\ddagger}$ | 15.9/ 21.4 | 17.0 / 21.7 | 16.9/ 20.2 |
| No. atoms | | | |
| Protein | 1954 | 1920 | 1028 |
| Ion | 1 | 1 | 0 |
| Water | 220 | 268 | 157 |
| R.m.s. deviations from ideal | | | |
| Bond lengths (Å) | 0.006 | 0.006 | 0.007 |
| Bond angles (°) | 0.85 | 0.81 | 0.91 |
| Ramachandran plot | | | |
| Favored (%) | 98.0 | 96.8 | 98.4 |
| Outliers (%) | 0 | 0 | 0 |
| PDB code | 6rix | 6riy | 6riz |

The highest resolution shell is shown in parentheses.

$^{\dagger}R_{merge} = \sum_h \sum_i | I_i - \langle I \rangle | / \sum_h \sum_I I_i$, where $I_i$ is the observed intensity of the *i*-th measurement of reflection h, and $\langle I \rangle$ is the average intensity of that reflection obtained from multiple observations.

$^{\ddagger}R = \sum||Fo|-|Fc||/\sum|Fo|$, where Fo and Fc are the observed and calculated structure factors, respectively, calculated for all data. $R_{free}$ was defined in Brünger, 1992[61].

$^{\&}CC_{1/2}$ was defined in Karplus et al., 2012[62].
