## Supplemental Table 2 for "Convergent evolution of the Hedgehog/Intein fold in protein splicing"

**Supplemental Table S2** Structural homology identified by DALI server.

| Protein | PDB | rmsd | Z-score | Length | # of residues | Seq. identity (%) |
|---|---|---|---|---|---|---|
| Translation Initiation Factor 5 (IF-5) | 1BKB | 2.3 | 2.7 | 49 | 136 | 16 |
| Eukaryotic Translation Initiation factor 5A2 (IF-5A2) | 3HKS | 2.1 | 2.6 | 49 | 142 | 16 |
| Elongation Factor P (EF-P) | 1UEB | 2.2 | 2.5 | 44 | 184 | 11 |
| PI-*Sce*I | 1DFA | 2.2 | 8.9 | 124 | 429 | 19 |
| 17-hedgehog | 1AT0 | 1.9 | 17.0 | 128 | 145 | 16 |
