## Supplemental Fig. 1 for "Convergent evolution of the Hedgehog/Intein fold in protein splicing"

**a**

| | Block A | Block B | Block F | Block G |
|---|---|---|---|---|
| *MP-Be*DnaB | Q/PLALNTEVPTPSG | GTEITASASHGWTT | PVKCIGIDTEDHLFQ | SRILTHN/T |
| *Bvi*IcmO | F/PQPLHSLVRMADG | GRSVEAARVHHWPV | PARCLVVADEMHCYI | HDIVTHN/C |
| *Dra*Snf2 | K/AQPLDAKVLTPLG | GASVEADAEHLWNV | PAQCIAVDAPDHLYV | GYIVTHN/T |
| *Mle*DnaB | K/ALALDTPLPTPTG | GTVIVADAQHQWPT | PVRCVEVDNAAHLYL | GMVPTHN/S |
| *Msm*DnaB1 | K/ALALDTPLPTPSG | GTAIVADAQHQWPT | PVRCVEVDNPEHLYL | GMVPTHN/S |
| *Mch*DnaB1 | K/ALALYTPLPTPSG | GTVIVADAAHQWPT | PVRCVEVDNPAHLYL | GMVPTHN/S |
| *Npu*DnaE | Y/CLSYETEILTVEY | GSVIRATSDHRFLT | NVYDIGVER-DHNFA | NGFIASN/C |
| *Npu*DnaB | G/CLAGDSLVTLVDS | GRKIRATGNHKFLT | EVFDLTVPG-LHNFV | NNIIVHN/S |
| gp41-1 | Y/CLDLKTQVQTPQG | GKEIICSEEHLFPT | ELIDIEVSG-NHLFY | NDILTHN/S |

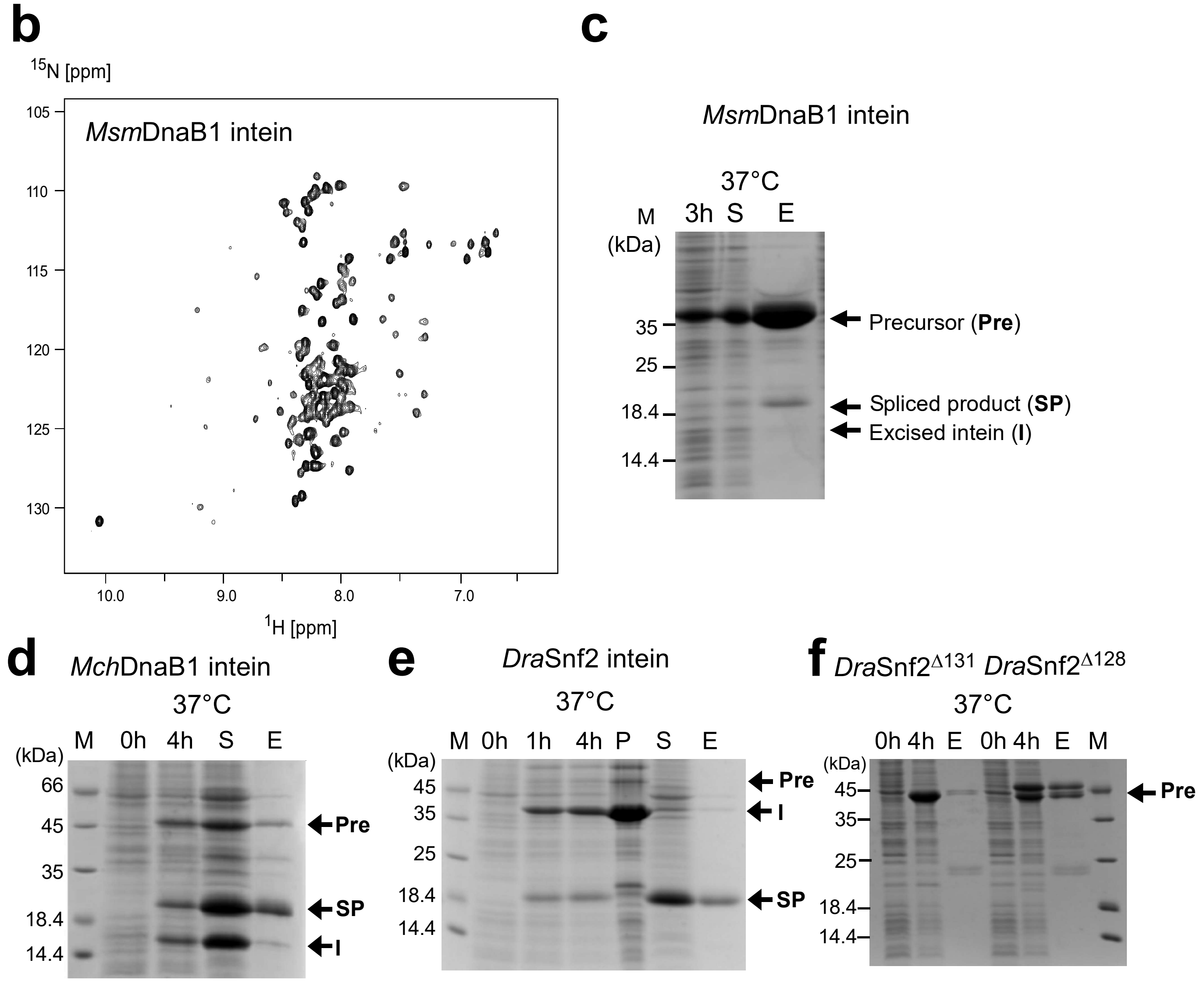


**Supplemental Figure S1:** Characterization of class 3 inteins. (**a**) A sequence alignment of various class 3 inteins (DnaB intein from *Mycobacteriophage Bethlehem* (MP-BeDnaB), IcmO intein from *Burkholderia vietnamiensis* G4 (BviIcmO), Snf2 intein from *Deinococcus radiodurans* (*Dra*Snf2), DnaB intein from *Mycobacterium leprae* (*Mle*DnaB), DnaB1 intein from Mycobacterium smegmatis (MsmDnaB1), and DnaB1 intein from *Mycobacterium chimaera* (*Mch*DnaB1)) and class 1 inteins (DnaE and DnaB inteins from *Nostoc punctiforme* (*Npu*DnaE, *Npu*DnaB) and gp41-1 intein). The alignment for blocks A, B, F, and G is shown. The WCT motif and the first residue are highlighted in yellow. (**b**) [$^{1}$H,$^{15}$N]-HSQC NMR spectrum of *Msm*DnaB1 intein, indicating poor folding of *Msm*DnaB1 intein. (**c-d**) SDS-PAGE analysis of protein splicing in *cis* by selected class 3 inteins, including two deletion variants of DraSnf2 missing the endonuclease domain (*Dra*Snf2$^{\Delta131}$ and *Dra*Snf2$^{\Delta128}$) (names indicated above).
