## Supplemental Fig. 2 for "Convergent evolution of the Hedgehog/Intein fold in protein splicing"

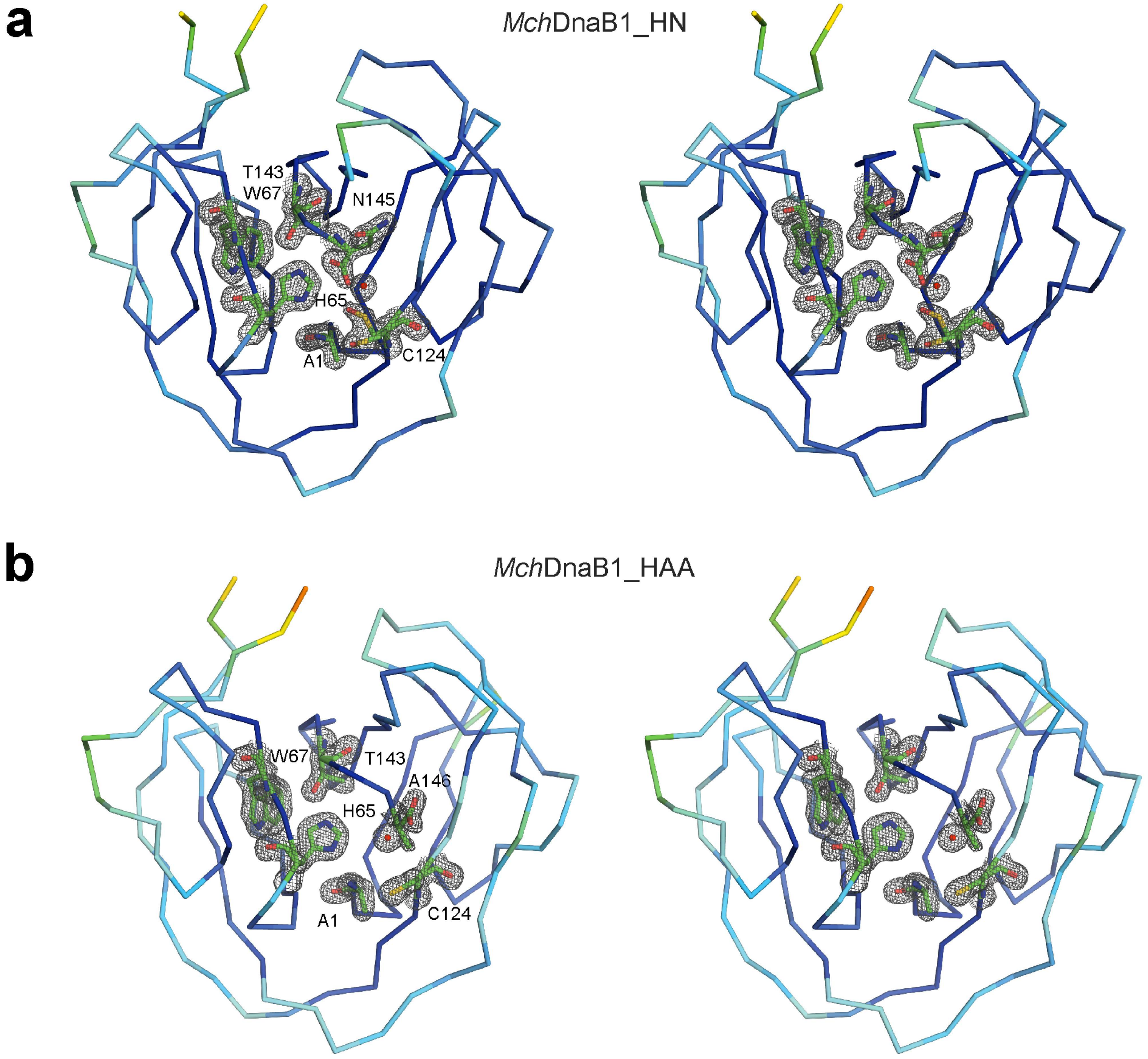


**Supplemental Figure S2:** The crystal structures of the *Mch*DnaB1 intein variants. Stereo-views of the backbone structures of (**a**) *Mch*DnaB1_HN and (**b**) *Mch*DnaB1_HAA showing the electron densities at the active-sites. The structures were depicted and colored according to the temperature factor using PyMol.
