## Supplemental Fig. 3 for "Convergent evolution of the Hedgehog/Intein fold in protein splicing"

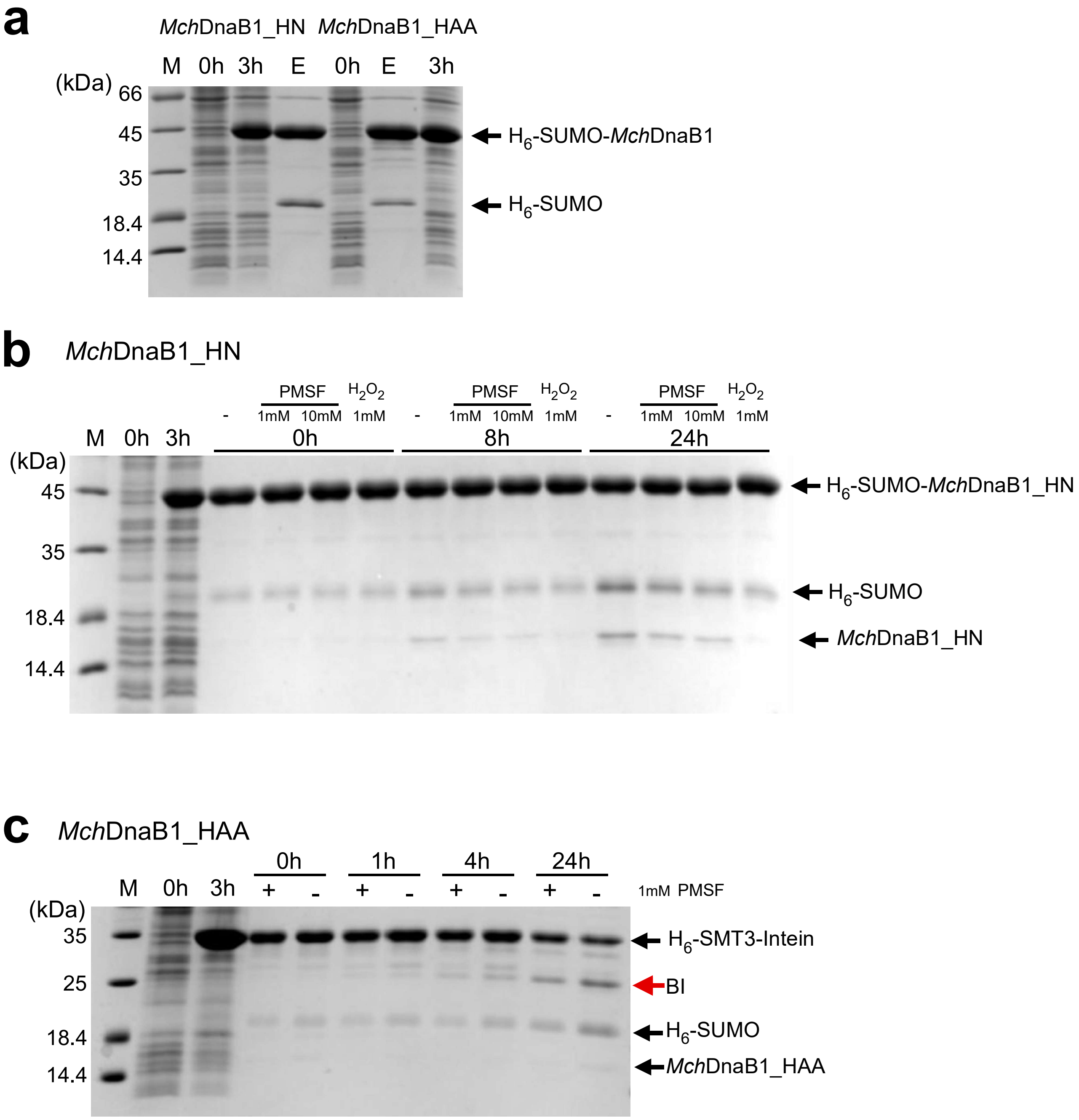


**Supplemental Figure S3**: N-cleavage of class3 *Mch*DnaB1 intein variants. (**a**) N-cleavage of MchDnaB1_HN and *Mch*DnaB1_HAA inteins immediately after purification. (**b-c**) Inhibition of N-cleavage of *Mch*DnaB1_HN (b) and *Mch*DnaB1_HAA (**c**) by addition of the indicated concentrartions of PMSF and $H_2O_2$ over time. (-), without inhibitor. After IMAC purification, the eluted protein was immediately incubated with the indicated inhibitors, and N-cleavage was monitored for 24 hours. Samples were analyzed by SDS-PAGE at the indicated time points. (**a-c**) Arrows indicate the corresponding bands for $H_6$-SUMO-intein, the precursor before cleavage, cleaved intein (*Mch*DnaB1_HN or *Mch*DnaB1_HAA), and cleaved $H_6$-SUMO. BI stands for the branched intermediate. M, 0h, and 3h indicate molecular weight marker, the sample before induction, and 3 hours after protein induction, respectively.
