## Supplemental Fig. 4 for "Convergent evolution of the Hedgehog/Intein fold in protein splicing"

**N-cleavage by salt-inducible class 1 intein ($H_6$-GB1-*Hut*MCM2_HAA-GB1)**

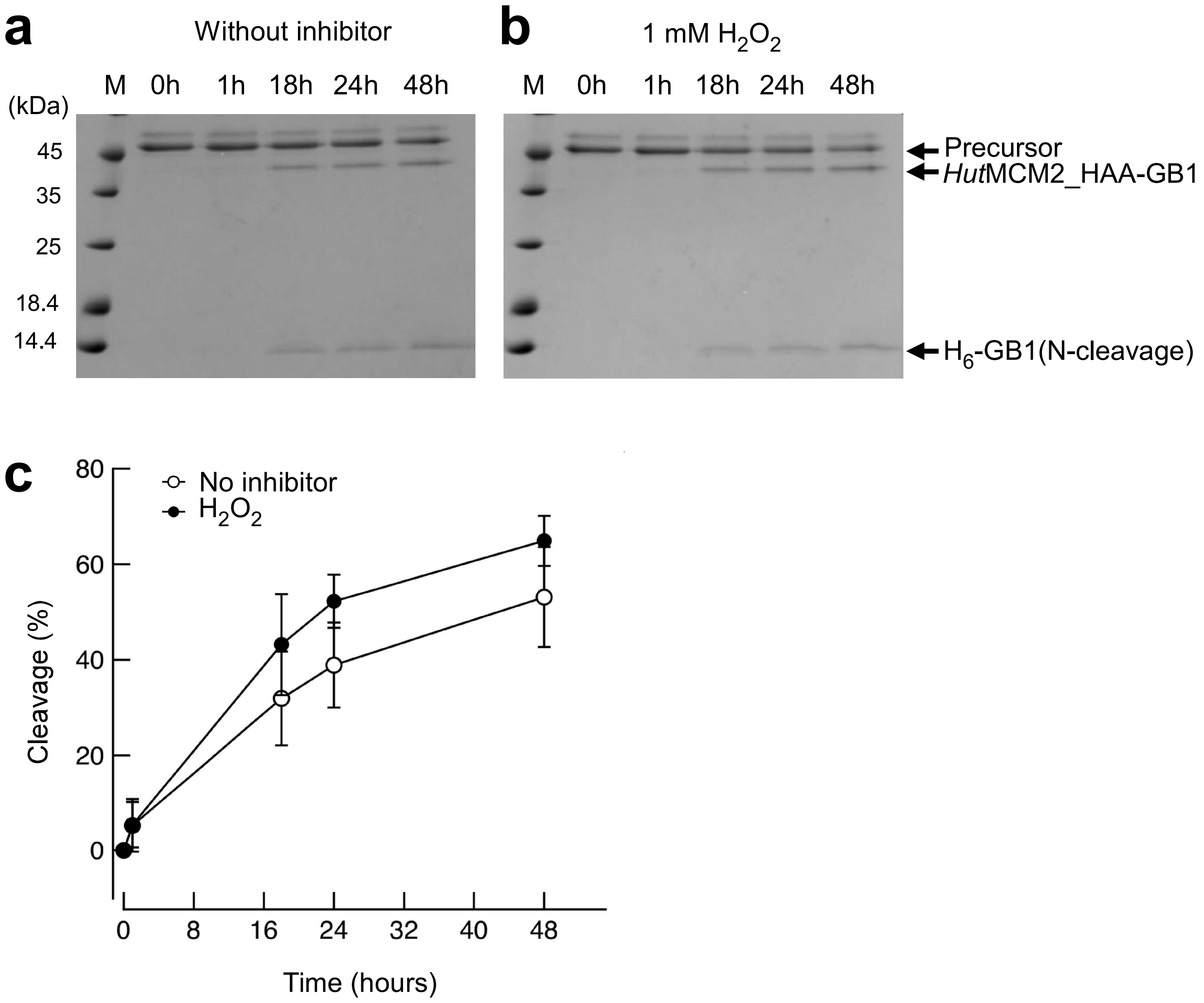


**Supplemental Figure S**4: Inhibition of N-cleavage of the class-1*Hut*MCM2_HAA intein by $H_2O_2$. After IMAC purification, the eluted protein was immediately incubated at high salinity (3.5 M NaCl) in the absence (**a**) or presence (**b**) of 1 mM $H_2O_2$ and N-cleavage was monitored for 48 hours by SDS-PAGE analysis. (**c**) Quantification of N-cleavage of the experiment in (**a**) and (**b**). Data were averaged from three individual experiments. Error bars represent one standard deviation.
