## Supplemental Fig. 5 for "Convergent evolution of the Hedgehog/Intein fold in protein splicing"

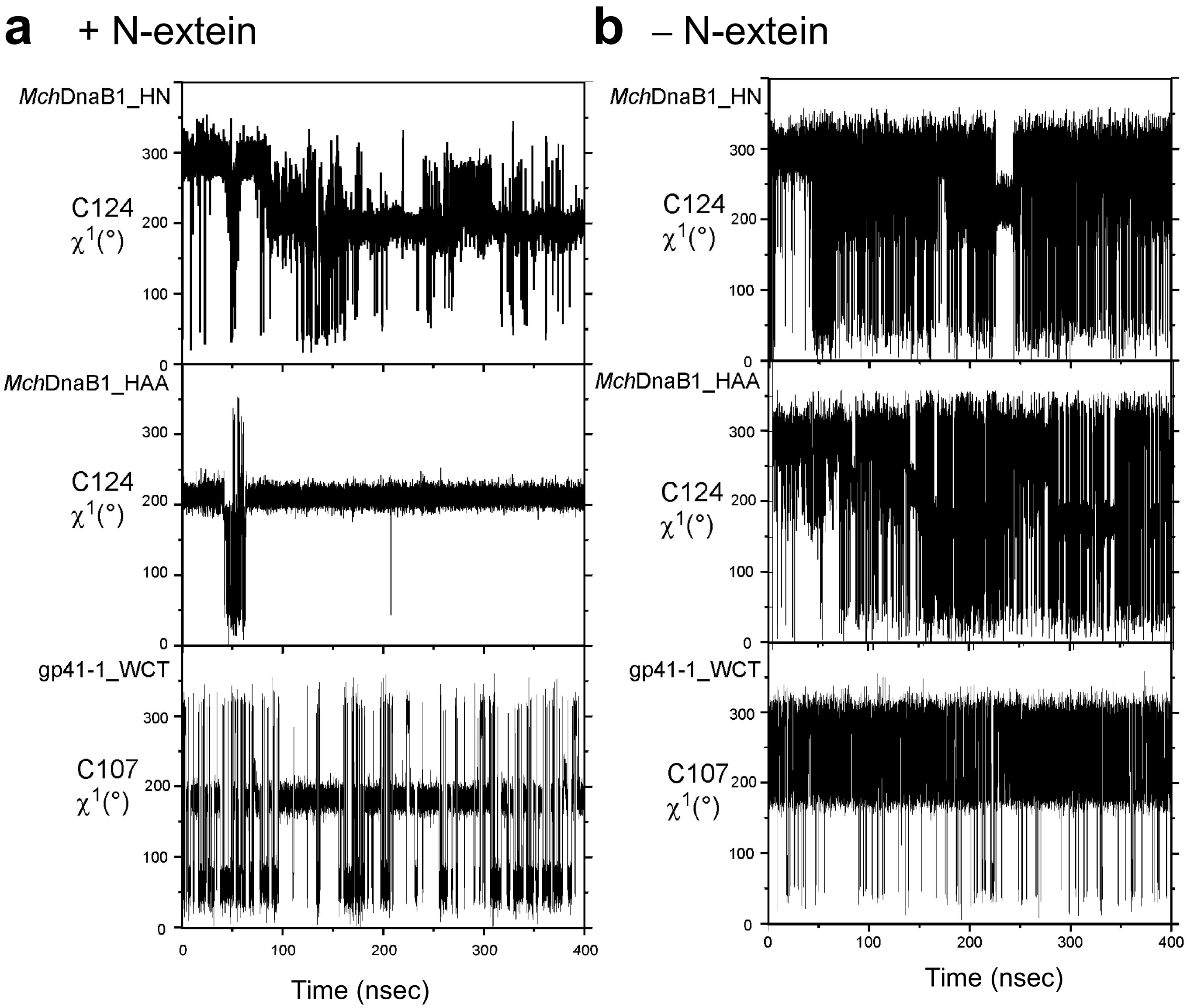


**Supplemental Figure S5**: MD simulations of the two variants of the *Mch*DnaB1 intein (*Mch*DnaB1_HN and *Mch*DnaB1_HAA) and the engineered gp41-1 intein with WCT motif (gp41-1_WCT). (**a**) Trajectories of the $\chi^1$ angle for the cysteine residues in the WCT motif during the 400-nano sec MD simulation with the modeled N-extein sequence. (**b**) Trajectories of the same $\chi^1$ angle without N-extein.
