## Supplemental Fig. 6 for "Convergent evolution of the Hedgehog/Intein fold in protein splicing"

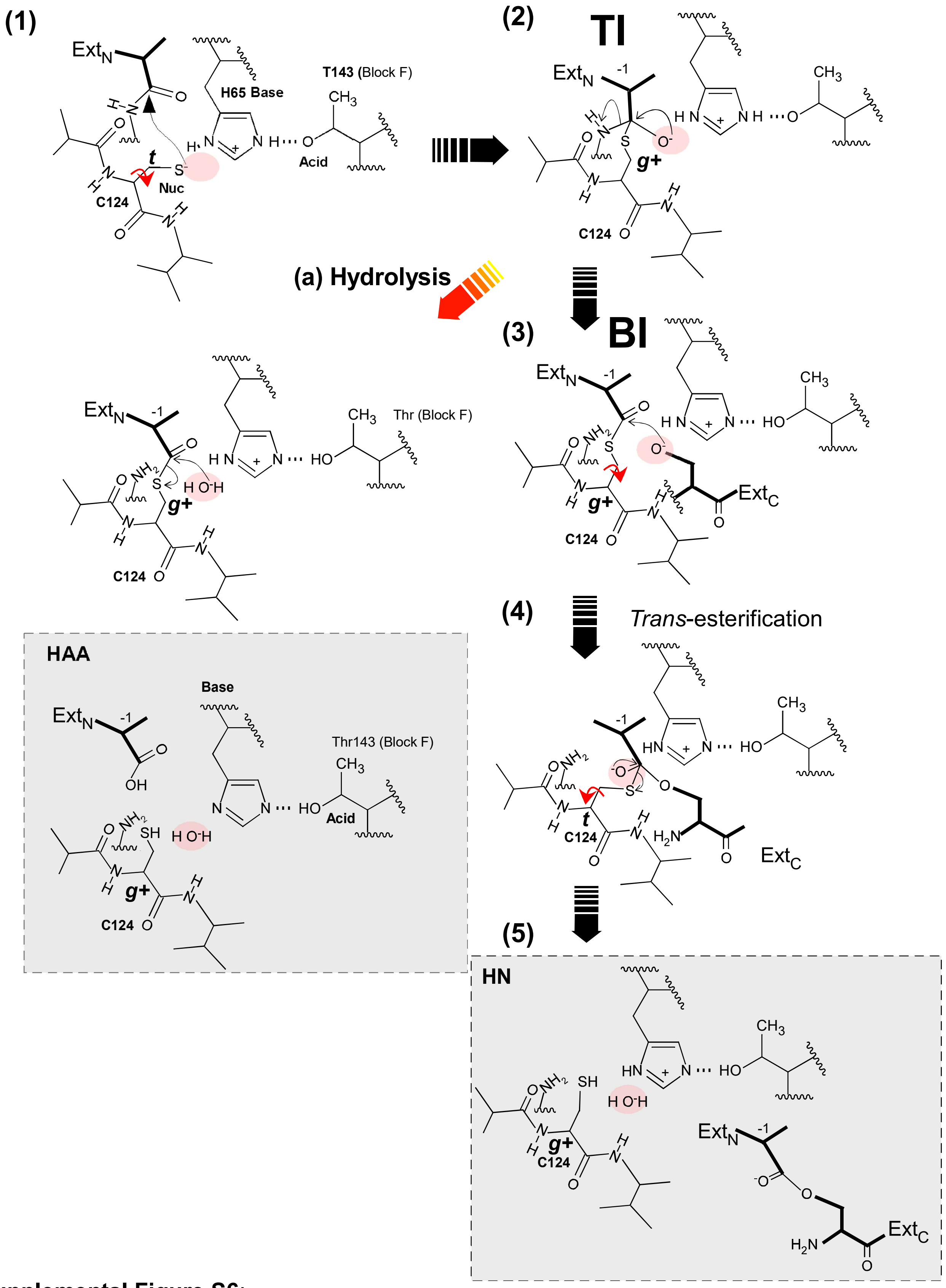


**Supplemental Figure S6**:
Proposed reaction steps for the protein splicing mechanism catalyzed by the class 3 intein and the relations to the solved crystal structures. **(1)** High energy ground state before splicing. **(2)** Tetrahedral Intermediate (**TI**) status after rotation of Cys124 to the gauche$^{+}$ conformation. **(3)** Branched Intermediate (**BI**) status. Rotation of Cys124 to the *trans* conformation will bring the thioester intermediate closer to the nucleophilic residue of the C-extein. **(4)** *Trans*-esterification step via a tetrahedral intermediate. Rotation of Cys124 back to the *gauche*+ conformation. (5) Post-splicing status. Exteins are released from the intein. HN and HAA stand for the conformations represented by the crystal structures of *Mch*DnaB1_HN and *Mch*DnaB1_HAA, respectively. (a) The non-productive N-cleavage off-pathway (hydrolysis) is indicated by a red arrow.
