## Supplemental Fig. 7 for "Convergent evolution of the Hedgehog/Intein fold in protein splicing"

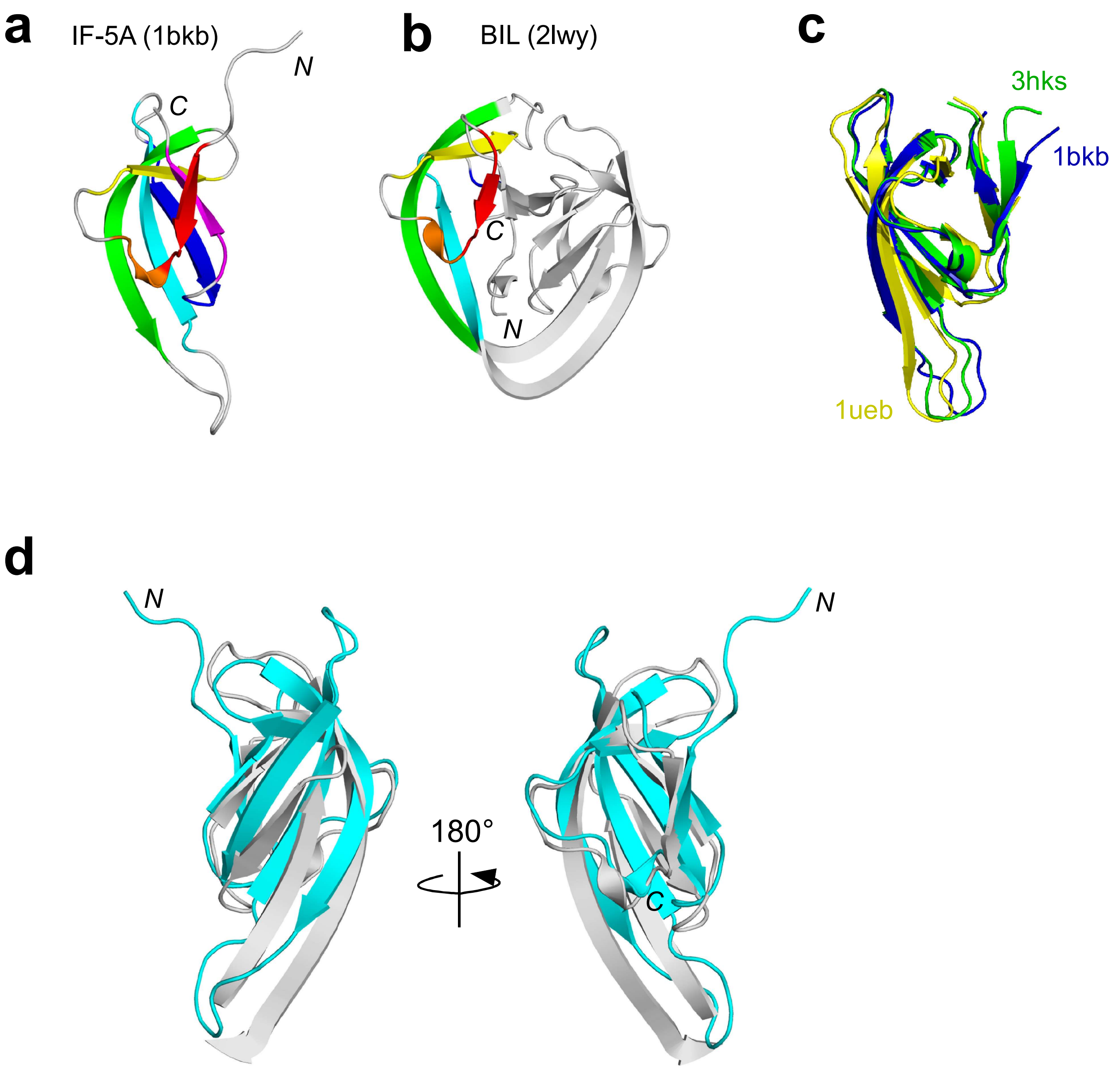


**Supplemental Figure S7**: Possible ancestral domains of the HINT fold. (**a**) The N-terminal domain of IF-5A from *Pyrobaculum aerophilum* in the crystal structure (1bkb). The secondary structures used for the structural alignment are colored. (**b**) The crystal structure of BIL4 domain from *Clostridium thermocellum* (2lwy). The secondary structures used for the superposition with the structure of IF-5A are colored with the same colors as in (**a**). (**c**) A superposition of the three crystal structures of Translation Initiation Factor 5 (IF-5A) from *Pyrobaculum aerophilum* (1bkb), Eukaryotic Translation Initiation factor 5A2 (3hks), and Elongation Factor P (1eub). (**d**) Overlays of the two structures from the N-terminal domain of IF-5A and *Cth*BIL4 domain. Only the regions of *Cth*BIL4 used for the superposition with the N-terminal domain of IF-5A are shown.
